# Adventitial macrophage accumulation impairs perivascular nerve function in mesenteric arteries with inflammatory bowel disease

**DOI:** 10.1101/2023.04.04.535591

**Authors:** Elizabeth A. Grunz, Benjamin W. Jones, Olubodun Lateef, Sidharth Sen, Katie Wilkenson, Trupti Joshi, Erika M. Boerman

## Abstract

*Introduction:* Inflammatory bowel disease (IBD) involves aberrant immune responses and is associated with both cardiovascular disease risk and altered intestinal blood flow. However, little is known about how IBD affects regulation of perivascular nerves that mediate blood flow. Previous work found perivascular nerve function is impaired in mesenteric arteries with IBD. The purpose of this study was to determine the mechanism of impaired perivascular nerve function.

*Methods:* RNA sequencing was performed on mesenteric arteries from IL10^-/-^ mice treated with *H.hepaticus* to induce disease (IBD) or left non-gavaged (Control). For all other studies, Control and IBD mice received either saline or clodronate liposome injections to study the effect of macrophage depletion. Perivascular nerve function was assessed using pressure myography and electrical field stimulation. Leukocyte populations, and perivascular nerves, and adventitial neurotransmitter receptors were labeled using fluorescent immunolabeling.

*Results:* IBD was associated with increased in macrophage-associated gene expression, and immunolabeling showed accumulation of adventitial macrophages. Clodronate liposome injection eliminated adventitial macrophages, which reversed significant attenuation of sensory vasodilation, sympathetic vasoconstriction and sensory inhibition of sympathetic constriction in IBD. Acetylcholine-mediated dilation was impaired in IBD and restored after macrophage depletion, but sensory dilation remained nitric oxide independent regardless of disease and/or macrophage presence.

*Conclusion:* Altered neuro-immune signaling between macrophages and perivascular nerves in the arterial adventitia contributes to impaired vasodilation, particularly via dilatory sensory nerves. Targeting the adventitial macrophage population may help preserve intestinal blood flow in IBD patients.

## 2. Introduction

Inflammatory Bowel Diseases, such as Crohn’s disease and ulcerative colitis, affect >3 million adults in the US, and prevalence increased 123% from 2007-2017(1). IBD is marked by chronic aberrant immune responses and both intestinal and extraintestinal inflammation. Patients experience increased risk and incidence of cardiovascular diseases including atherosclerosis(2), stroke(3), myocardial infarction(4) and heart failure(5) despite having lower traditional risk factors (obesity, hypertension, hyperlipidemia, diabetes) than the general population(4). IBD decreases intestinal perfusion and increases bowel ischemia (6, 7, 8), and blood flow measurements are used in diagnosis and assessment of relapse risk(9). Collectively, these factors point to an important role for vascular function in the pathogenesis and treatment of IBD, yet few studies have directly addressed changes in vascular function with IBD.

Mesenteric arteries (MAs) direct blood to the intestines, and their function is critical to mobilizing ∼25% of cardiac output in and out of the gut(10). The perivascular sympathetic and sensory nerves enmeshed in the adventitia directly constrict and dilate MAs, respectively, (11) to regulate perfusion. Perivascular sensory nerves further facilitate dilation via feedback inhibition of sympathetic nerves(12, 13) and local axon reflexes(14). Despite diminished vasodilator capacity(15) and blood flow(6, 8, 16) with IBD, the function of sensory nerves and their neuroeffector signaling pathways in IBD is understudied. Previous work in the lab showed significant defects in both sensory vasodilation and sensory inhibition of sympathetic vasoconstriction with IBD (17). These deficits coincide with modest changes in sensory neurotransmitter content, release and receptor function, but the specific mechanism underlying the near-total loss of sensory vasodilation in IBD remains undefined.

The adventitia that houses perivascular nerves contains a dynamic mix of perivascular nerves, immune cells, pericytes, and extracellular matrix that participate in paracrine signaling in both normal and pathological function(18, 19). In healthy adventitia, immune cells are relatively quiescent, and sympathetic and sensory nerve activity are balanced by reciprocal feedback(12, 13). Many cardiovascular and inflammatory diseases are accompanied by an increase in the number and/or change in phenotype of adventitial(20) immune cells, particularly macrophages. When activated by local signals, macrophages polarize into functional subpopulations across a spectrum including classically activated M1-like, and alternatively activated M2-like phenotypes(21). In IBD, macrophage accumulation and subtype disequilibrium in the colon wall contributes to disease pathogenesis and severity(22, 23) where decreased blood flow is evident, but no studies have investigated adventitial macrophages with respect to vascular function or blood flow in IBD. The role of perivascular sensory nerves in these processes is also unclear. Using RNA sequencing, this study identified increased macrophage markers in arteries with IBD and therefore tested whether macrophages and other leukocytes accumulate in the mesenteric artery adventitia in the presence of established IBD. Given our previous findings of perivascular nerve dysfunction (17), we therefore tested whether and how whole-body macrophage depletion improves perivascular nerve function.

## 3. Materials and Methods

### 2.1 Animals and Artery Preparation

All experiments in the present study utilized the widely-used and accepted IL10^-/-^ model of IBD, where IBD was induced by administration of the commensal gut bacteria *Helicobacter hepaticus*. Briefly, male and female IL-10^−/−^ mice (4–6Cmo), originally obtained from the Jackson Laboratory (B6.129P2-IL-10tm1Cgn/J, Strain #002251, RRID:IMSR_JAX:002251) were bred and housed at the University of Missouri in a 12:12 light:dark cycle and received standard chow and water ad libitum. At 2 and 4 days post-weaning, mice in the IBD group were administered *H. hepaticus* by oral gavage as previously described(17) and developed disease for 90 days as shown in **Figure 1A**. For studies using macrophage depletion, Control and IBD groups received intraperitoneal injections of PBS- or clodronate liposomes (100 µl, 50 µl and 50 µl of 5mg/ml stock in PBS) at days 111, 118 and 125, respectively, as shown in **Figure 1B**. For all experiments, mice were anesthetized via intraperitoneal injection of ketamine-xylazine (25 mg/mL/2.5 mg/mL) followed by tissue dissection and euthanasia via cardiac exsanguination. All experiments were performed in compliance with the Guide for the Care and Use of Laboratory Animals (24) and were approved by the University of Missouri Animal Care and Use Committee. Inflammation of the colon was determined post-hoc via blind analysis in H&E-stained intestine sections using a previously-described scoring system (17, 25) ranging from 0-24. For all experiments, first or second-order mesenteric arteries (MAs, 100-200 µm inner diameter) from the terminal ileum were isolated by hand dissection in Ca^2+^ free physiological salt solution (PSS) [in mM: 140 NaCl, 5 KCl, 1 MgCl2, 10 HEPES, 10 glucose] at 4CC.

**Figure 1.**
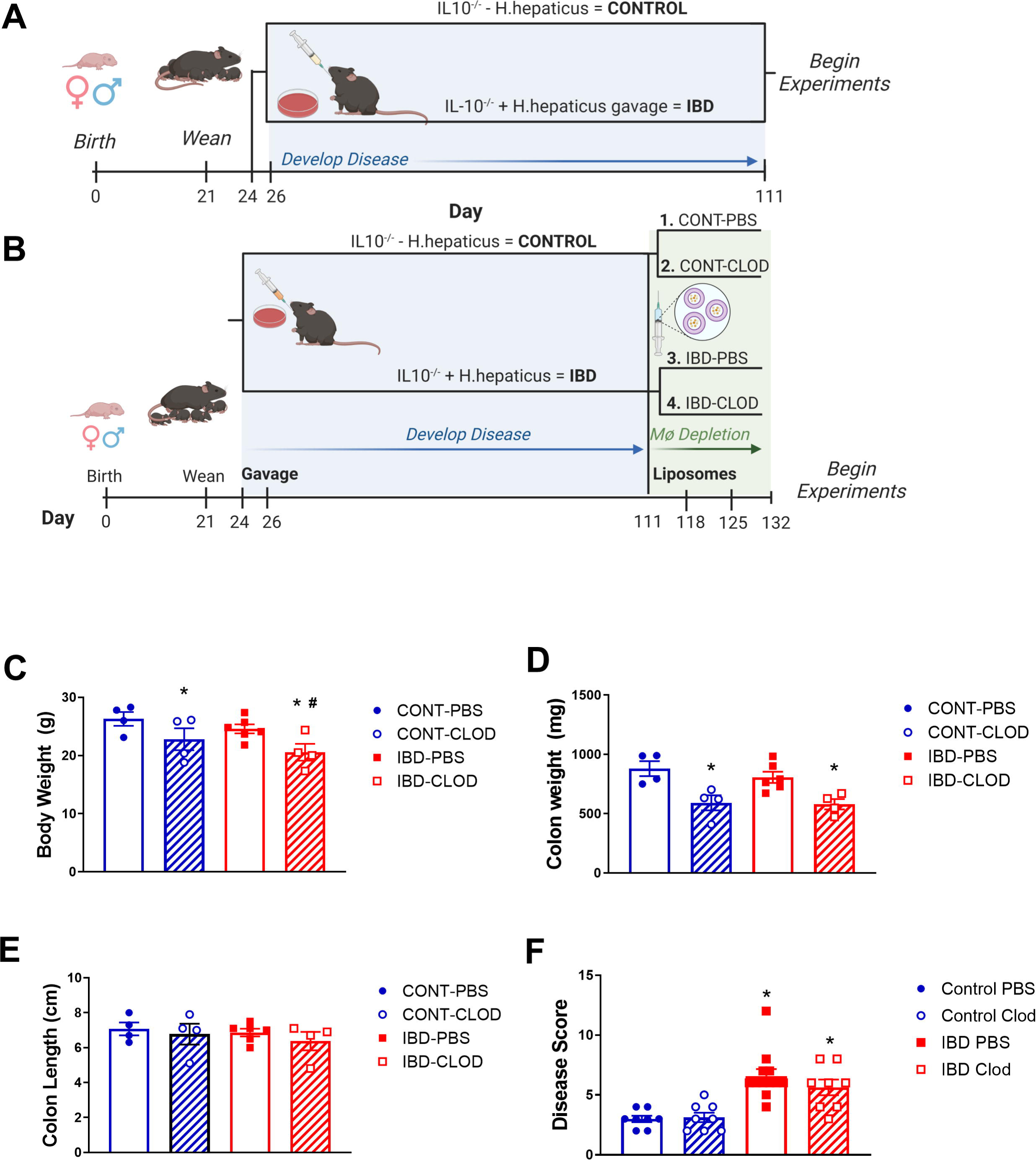
Experimental IBD models and associated mouse data. Figures depict experimental timelines for studies A) without macrophage depletion and B) with macrophage depletion via clodronate liposome injection. Note that in all experiments, Control animals are non-gavaged IL10^-/-^ mice, while IBD groups are IL10^-/-^ mice gavaged with *H. hepaticus* after weaning. For all groups measurements were collected for mean C) body weight, D) colon weight, E) colon length and F) histological disease score of H&E stained colon sections in Control (CONT) and IBD mice treated with either PBS or clodronate (CLOD) liposomes. N=4-8 mice per measurement in each group. * = P<0.05 vs CONT-PBS; # = P<0.05 vs IBD-PBS. Timelines created in Biorender.

### 2.2 RNA sequencing

First and second-order mesenteric arteries from each mouse were pooled and placed into lysis buffer from the RNAqueous Micro Kit with DNase from Ambion Inc. (Austin, TX, USA), followed by RNA extraction and DNase treatment per manufacturer’s guidelines. All samples were provided to the DNA Core at the University of Missouri for RNA sequencing. Libraries were prepared using the TruSeq Stranded mRNA Library Kit according to the manufacturer’s instructions, with quantity and quality verified using the High Sensitivity NGS Fragment Analysis Kit (Agilent Technologies). Libraries were pooled in equimolar amounts and sequenced with single end 75 base pair reads using an Illumina NextSeq 550. Resulting Bcl files were converted to Fastq format using Illumina software, and trimming was performed using Trimomatic (26) with a threshold of 36. Reads were aligned to the GRCm38.p5 Mus musculus annotated Genome assembly from Ensembl. Transcript abundance (expressed in FPKM) was generated using the Cufflinks tool from the Tuxedo suite (27) as well as genome annotation files downloaded from the same source. Differential gene expression analysis was performed using the Cuffdiff tool(27). Differentially expressed transcripts were defined as those that had a fold change with an adjusted P < 0.05 via Wald test and by Benjamini–Hochberg correction. Clustering, principal component analysis (PCA) and heatmap generation were performed on ln(x)-transformed data using ClustVis software(28). The heatmap was generated using row centering via unit variance scaling. Rows and columns were clustered using correlation distance and average linkage. Principal component analysis used singular value decomposition with imputation. Relevant GO terms were determined based on the list of differentially expressed genes using the GO Enrichment Analyzer web tool (http://geneontology.org/).

### 2.3 Pressure Myography

Unbranched Second-order arteries 500-1000 µm in length were transferred to a cannulation chamber with platinum electrodes (Warner Instruments), cannulated onto glass pipettes, and tied with two 11-0 sutures on each end. After transferring the rig to microscope stage, arteries were pressurized to 70CmmHg and heated to 37°C. For all functional studies, arteries were superfused with PSS (above) containing 2 mM calcium. Images were acquired on an inverted Olympus BX61W1 microscope using a 10X objective and a CCD camera (Basler, Inc) coupled to edgeCtracking software (LabView, National Instruments). Perivascular nerves were stimulated in an electrical field as previously described (17). Briefly, electrical pulses (70 V, 2 ms) were delivered using a Grass S88 stimulator and stimulus isolation unit at 1-16 Hz until a stable diameter was achieved (<30 s). Sympathetic constrictions were measured in the presence and absence of capsaicin (10 μM), which blocks sensory nerves. Sensory vasodilation was measured in vessels preconstricted with phenylephrine (1 μM) and the sympathetic nerve blocker guanethidine (10 μM). To determine the endothelium dependence of sensory vasodilation, stimulations were repeated in the presence of the nitric oxide inhibitor N(G)-Nitro-L-arginine methyl ester (L-NAME, 100 μM). In all experiments, maximum diameters were determined by incubating vessels in Ca^2+^ free PSS containing 10 μM sodium nitroprusside.

### 2.4 Immunofluorescence

Isolated mesenteric arteries cleared of perivascular adipose were pinned in a Sylgard-coated 24Cwell plate for immunolabeling. Arteries were fixed in 4% paraformaldehyde, blocked and permeabilized with phosphate buffered saline (PBS) containing 1% bovine serum albumin and 0.1% Triton XC100, and incubated overnight in primary antibodies (details in **Table 1**). Immune cells were labeled as follows: leukocytes with CD45, M1-like macrophages with CD68, M2-like macrophages with CD163, antigen presenting cells with MHCII. In separate experiments, total macrophages were labeled with F4/80, calcitonin gene-related peptide (CGRP) receptors with the receptor activity modifying protein 1 (RAMP1) and the neurokinin 1 receptor (NK1) for substance P. Perivascular nerves were labeled as follows: all nerves with PGP 9.5, sympathetic nerves with tyrosine hydroxylase, sensory nerves with CGRP and substance P. Tissues were then blocked again, incubated in secondary antibodies, and mounted on slides using ProLong Gold with or without DAPI. Each experiment included no primary controls that demonstrated a lack of nonspecific fluorescence. All slides were imaged on a Leica TCS SP8 laser scanning confocal microscope at 25X with 1 µm z-sections. Each fluorophore was imaged sequentially using a separate laser line for excitation to prevent cross-excitation. Representative images were prepared by generating maximum z-projections through one vessel wall. Immune cells were counted manually post-hoc. Nerve density was quantified from maximum projections in Image J (29) using a previously-described procedure (17) to measure the fluorescence area fraction of each antibody. CGRP and SP receptor (RAMP1 and NK1, respectively) colocalization with macrophage F4/80 labeling was measured using an object-based colocalization plugin in FIJI (30) that measures colocalization within z-stacks in both whole cells and their nuclei.

**Table 1.**
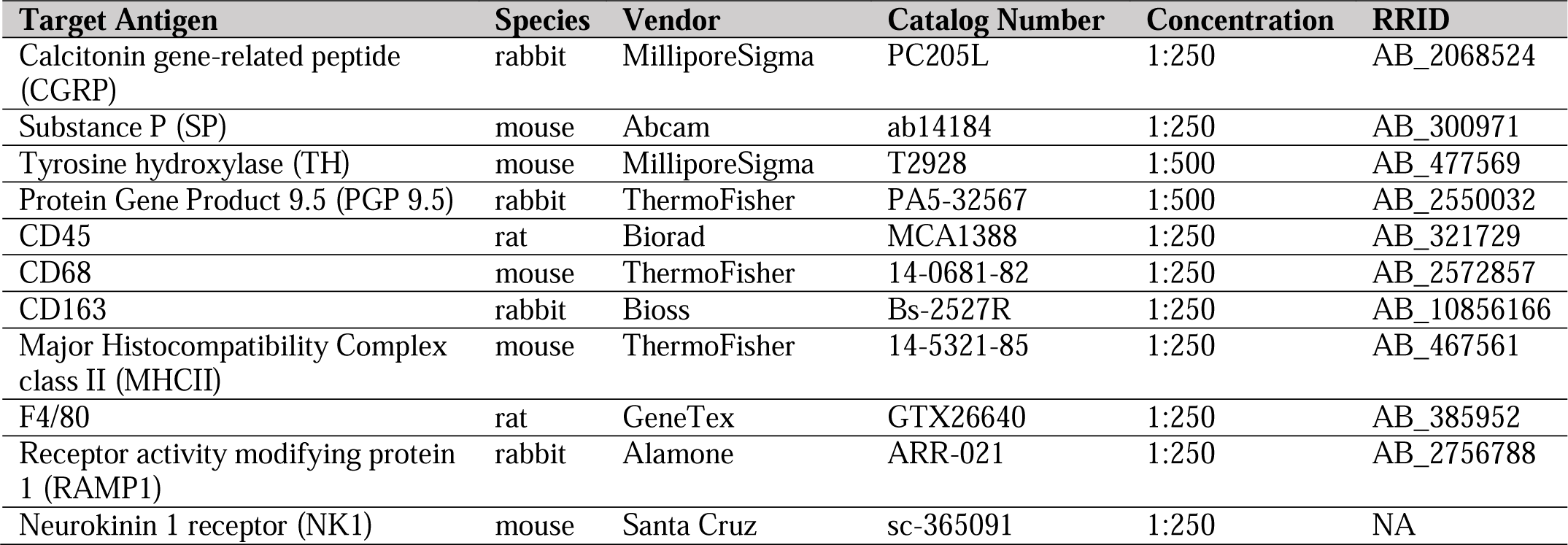
Primary antibody information.

### 2.5 Chemicals, Reagents and Tools

As described above, antibodies used in all immunolabeling experiments are outlined in **Table 1**. All other key reagents, chemicals and drugs are listed in **Table 2** with their vendor, catalog number, and concentration(s) used.

**Table 2.** Drug information.

| Name | Vendor | Catalog Number | Concentration | Vehicle |
| --- | --- | --- | --- | --- |
| Phenylephrine hydrochloride (PE) | MilliporeSigma | P6126 | 1 μM | Water |
| Guanethidine sulfate (GE) | BOC Sciences | 645-43-2 | 10 μM | Water |
| Sodium nitroferricyanide (nitroprusside, SNP) | MilliporeSigma | 228710 | 10 μM | Water |
| Capsaicin (CAP) | MilliporeSigma | M2028 | 10 μM | Ethanol |
| Nω-Nitro-L-arginine methyl ester hydrochloride (L-NAME) | MilliporeSigma | N5751 | 100 μM | Water |
| Clodronate liposomes | Liposoma | C-005 | 5mg/ml stock | PBS |
| PBS liposomes | Liposoma | P-005 | -- | PBS |

### 2.6 Statistical Analysis

All data are reported as means ± SEM, with significance at P < 0.05. n values represent the number of biological replicates per group used in each experiment unless otherwise noted. Statistical analysis was completed using Graphpad Prism 9. All concentration-response and electrical field stimulation data were analyzed by two-way repeated measures ANOVA followed by Bonferroni post hoc test, with differences tested within and between groups. Disease scoring, colon length and body weights were analyzed by unpaired t tests. Fluorescence imaging analysis for immune cell counts and nerve density were analyzed via nested t-test to account for the use of multiple images from each biological replicate.

## 4. Results

### 4.1 IBD induction and macrophage depletion in IL10^-/-^ mice

In the present study, all studies examined the effects of IBD using one of two experimental protocols. In the standard IBD protocol (**Figure 1A**), studies were conducted all mice 90 days post-gavage. Experiments examining the effect of macrophage depletion used the clodronate protocol (**Figure 1B**) in which mice were treated with either PBS liposomes (macrophages intact) or clodronate liposomes (macrophages depleted). In mice treated with PBS liposomes, there was a non-significant trend toward decreased body weight (**Figure 1C**), and no changes in colon weight (**Figure 1D**) or length (**Figure 1E**) 90 days post-gavage. Conversely, and consistent with our previous studies(17), IBD induction with *H. hepaticus* gavage was associated with a significant increase in histological disease scores IBD vs Control colons (**Figure 1F**) led to a significant increase in colon inflammation scores. Clodronate liposome treatment was associated with decreased body weight in Control and IBD groups, with a significantly greater decrease in IBD-CLOD mice (**Figure 1C**). Clodronate decreased colon weight to a similar extent in Control and IBD groups compared to PBS treatment (**Figure 1D**), but it had no effect on colon length (**Figure 1E**) or disease scores (**Figure 1F**) in either group compared to PBS treatment.

### 4.2 Increased expression of macrophage marker genes in mesenteric arteries with IBD

RNA sequencing of mesenteric arteries from identified a total of 228 genes with significantly differential expression in IBD vs Control, 152 with increased expression and 76 with decreased expression (**Supplemental Table 1**). Hierarchical heatmap creation also showed that individual Control and IBD samples segregated together based on differential gene expression patterns (**Figure 2A**). Principal component analysis determined that PC1 and PC2 combined account for 73.9% of variability in gene expression (**Figure 2A**). PC1 and PC2 were also used to look for any apparent patterns that clearly differentiate males from females in this group of genes, with none uncovered. The top 30 genes with upregulated expression in arteries from IBD mice included a number of macrophage marker genes including *cd163* (CD163), *cd68*, *itgam* (CD11b), *adgre1* (F4/80), *csf1r* (CD115) and *mrc1* (CD206). Additional genes found in the top 30 are not macrophage markers but associated with macrophage function: *ms4a4a*, *ccl9*, *siglec1* (CD169), *clec10a* (CD301). All significantly up- and downregulated genes are shown in in **Supplemental Table 1**, including the complete list of differentially expressed genes with fold changes, P and Q values. Consistent with an expanded macrophage population, GO enrichment analysis showed trends of enrichment of gene sets associated with immune cell chemotaxis and inflammation (**Figure 2C**). Based on these data, we conducted additional experiments looking at changes in the adventitial macrophage population in IBD and its role in perivascular nerve function/dysfunction.

**Figure 2.**
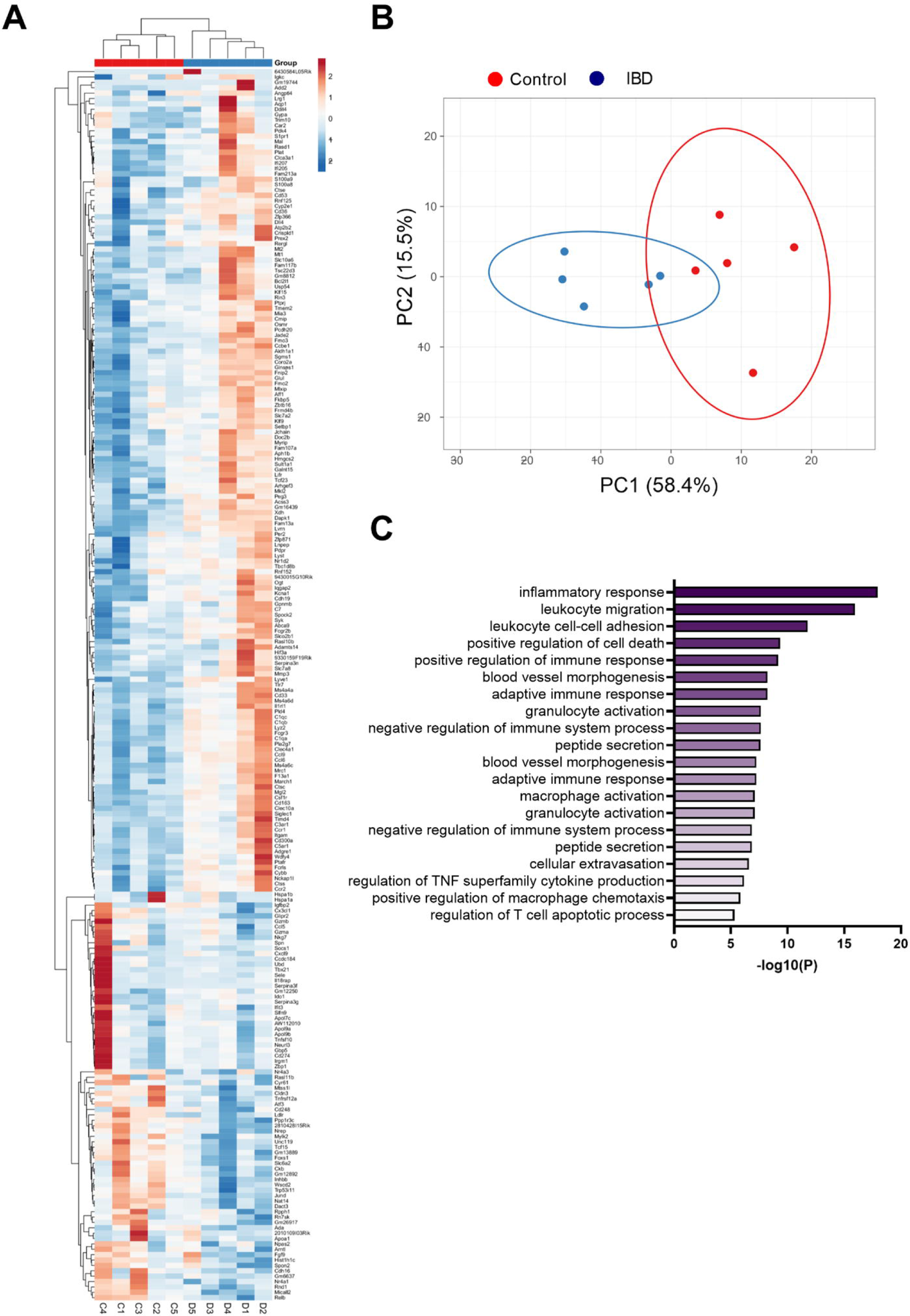
Heatmap, principal component analysis, and GO enrichment of differentially expressed genes from RNA sequencing of mesenteric arteries. A) Clustered heatmap of differentially expressed genes in mesenteric arteries from Control and IBD mice. Red colors indicate genes with increased expression in IBD vs Control, while blue colors represent genes with decreased expression. All differences have P < 0.05. B) Principal component analysis of differentially expressed genes from Control and IBD mice. C) Top 20 significantly enriched GO biological processes in mesenteric arteries from IBD vs Control mice. All data originated from RNA sequencing of N=5 samples each from Control and IBD groups. Note: a complete list of differentially expressed genes is found in **Supplemental Table 1**.

### 4.3 IBD increases, clodronate decreases adventitial immune cells

To characterize the adventitial leukocyte and macrophage populations in Control and IBD arteries with and without macrophage depletion, we conducted two sets of confocal immunolabeling studies counting the number of cell labeled for: CD45 (all leukocytes), MCHII (antigen presenting cells), CD68 (M1-like macrophages) and CD163 (M2-like macrophages). All markers were significantly increased in IBD-PBS arteries vs CONT-PBS arteries (**Figure 3**). Because CD163 antibodies exhibited some non-macrophage labeling, we confirmed the increase in M2 macrophages from IBD vs Control arteries (but not in macrophage-depleted mice) using a CD206 antibody, where quantitation showed a two-fold increase in CD206+ fluorescence area with IBD (P<0.05, data not shown). Macrophage depletion with clodronate led to significant attenuation of CD45+ cells in both CONT-CLOD and IBD-CLOD groups, although significantly more CD45+ cells persisted post-clodronate in CONT-CLOD. Clodronate treatment resulted in near-total elimination of MHCII+, CD68+ and CD163+ cells in both CONT-CLOD and IBD-CLOD groups, indicating successful depletion of macrophages in the artery adventitia (**Figures 3**). This is consistent with control flow cytometry experiments showing similar levels of macrophage depletion in the spleens of the same experimental mice (data not shown).

**Figure 3.**
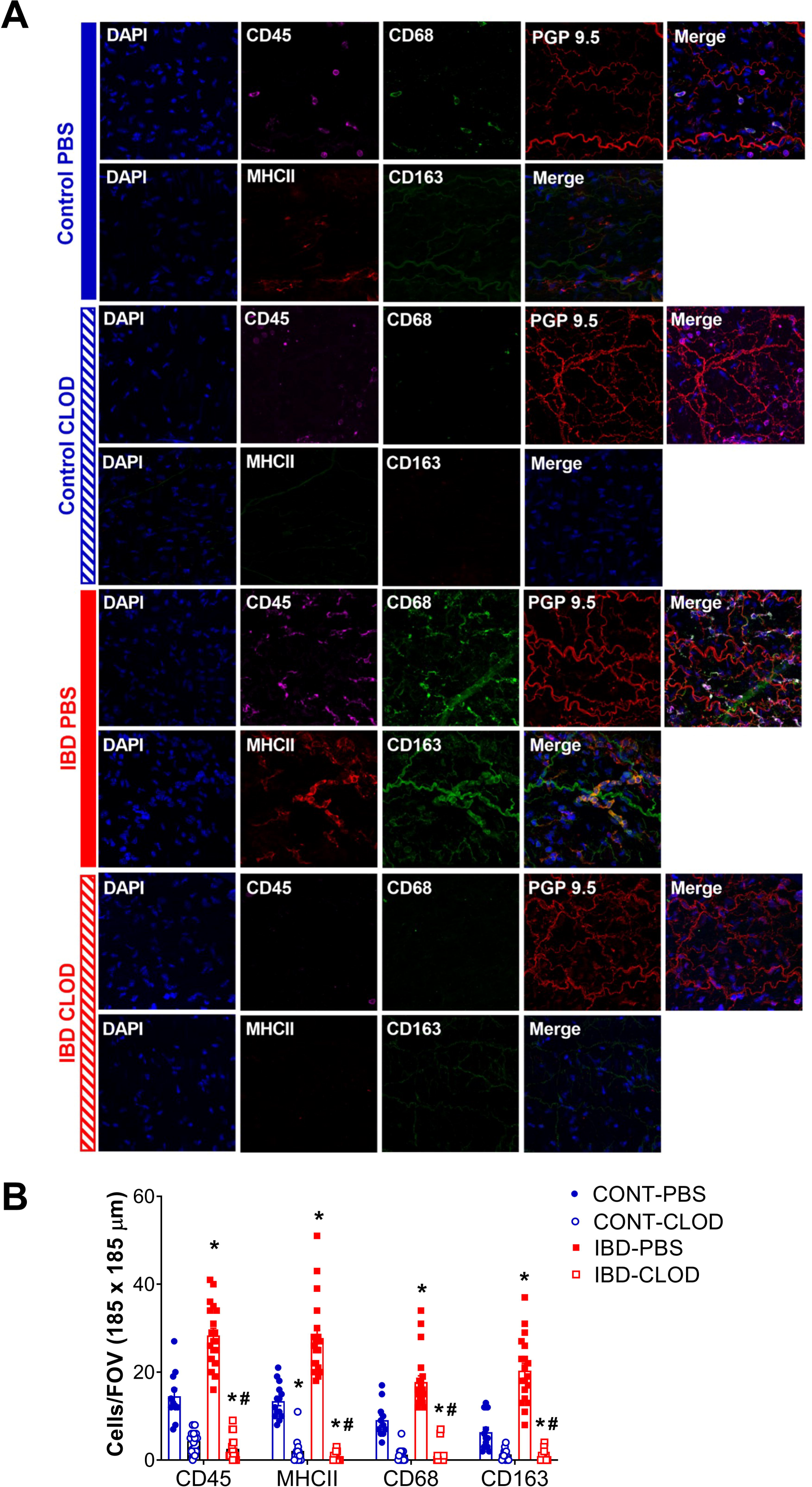
IBD increases and clodronate decreases adventitial macrophages and leukocytes. A) Images are representative maximum z-projections through one artery wall showing nuclei leukocyte, M1-like macrophages and perivascular nerves in mesenteric arteries in each of 4 groups: Control-PBS, Control-CLOD, IBD-PBS and IBD-CLOD. Panels for each group depict two separate image sets-Top row (left to right): Nuclei (DAPI, blue), leukocytes (CD45, magenta), M1-like macrophages (CD68, green), perivascular nerves (PGP9.5, red), merged image; Bottom row (left to right): nuclei (DAPI, blue), antigen presenting cells (MHC II, red), M2-like macrophages (CD163, green), merged image. Images are 185 × 185 µm. B) Quantitation of immunofluorescence for adventitial immune cells. Data are mean ± SE number of leukocytes (CD45+), antigen-presenting cells (MHCII+), M1-like macrophages (CD68+) and M2-like macrophages (CD68+) per field of view (FOV, 185 × 185 µm) of confocal z-projections through the mesenteric artery adventitia of vessels from Control and IBD mice treated with PBS or clodronate (CLOD) liposomes. N=12-20 images from 4 mice per group. * = P<0.05 vs Control-PBS; # = P<0.05 vs IBD-PBS.

### 4.4 Macrophage depletion improves perivascular nerve dysfunction in mesenteric arteries with IBD

#### 4.4.1 Macrophage depletion restores impaired sympathetic vasoconstriction and sensory inhibition of sympathetic vasodilation with IBD

Pressure myography and electrical field stimulation were used to assess sympathetic vasoconstriction in the presence (PSS) and absence (capsaicin) of sensory neurotransmission in CONT-PBS, CONT-CLOD, IBD-PBS and IBD-CLOD groups. Consistent with previous findings, sympathetic constriction was decreased in mesenteric arteries from IBD-PBS vs CONT-PBS mice. Despite impaired sympathetic constriction with IBD, the presence of sensory neurotransmission failed to decrease sympathetic vasoconstriction to even a small degree (**Figure 4A,C**). Macrophage depletion had opposing effects in Control and IBD groups. CONT-CLOD arteries exhibited decreased sympathetic vasoconstriction compared to CONT-PBS while retaining the ability of sensory nerves to oppose sympathetic vasoconstriction (i.e., capsaicin still increased sympathetic constriction in CONT-CLOD arteries) (**Figure 4A,B**). In contrast, macrophage depletion increased sympathetic constriction in IBD-CLOD vs IBD-PBS (**Figure 4C,D**), restoring them to CONT-PBS levels. Similarly, macrophage depletion restored sensory inhibition of sympathetic inhibition (**Figure 4C,D**), with IBD-CLOD constrictions in PSS and capsaicin becoming indistinguishable from CONT-PBS (**Figure 4A,D**).

**Figure 4.**
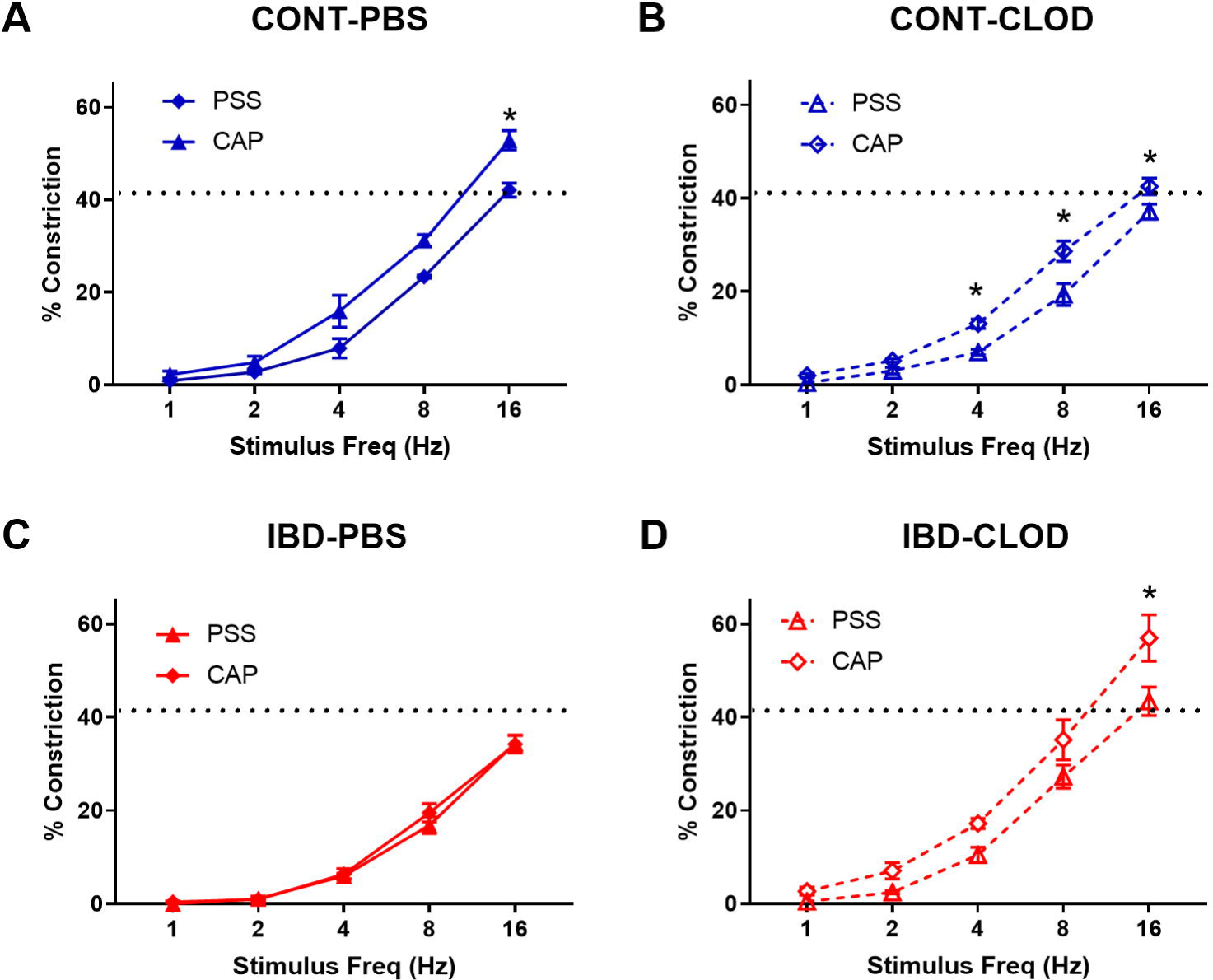
Macrophage depletion improves sensory inhibition of sympathetic vasodilation. Data are mean ± SE for %constrictions to electrical field stimulation (1-16 Hz) before (PSS) and after sensory nerve blockade with capsaicin (CAP, 10 µM) in A) CONT-PBS: Control mice treated with PBS liposomes, B) CONT-CLOD: Control mice treated with clodronate liposomes, C) IBD-PBS: IBD mice treated with PBS liposomes, and D) IBD-CLOD: IBD mice treated with clodronate liposomes. Dashed line indicates the max sympathetic constriction reached in CONT-PBS arteries in PSS, which was significantly greater vs IBD-PBS. N = 4-5 per group. * = P< 0.05 vs PSS

#### 4.4.2 Macrophage depletion restores impaired sensory vasodilation in mesenteric arteries with IBD

Consistent with previous studies in our group, we found that IBD was associated with significantly attenuated perivascular sensory vasodilation, with a ∼75% decrease in maximum achievable vasodilation in IBD-PBS vs CONT-PBS arteries (**Figure 5**). Macrophage depletion did not alter sensory vasodilation in CONT-CLOD arteries. In contrast, macrophage depletion increased sensory vasodilation in IBD-CLOD arteries, restoring dilations to Control levels.

**Figure 5.**
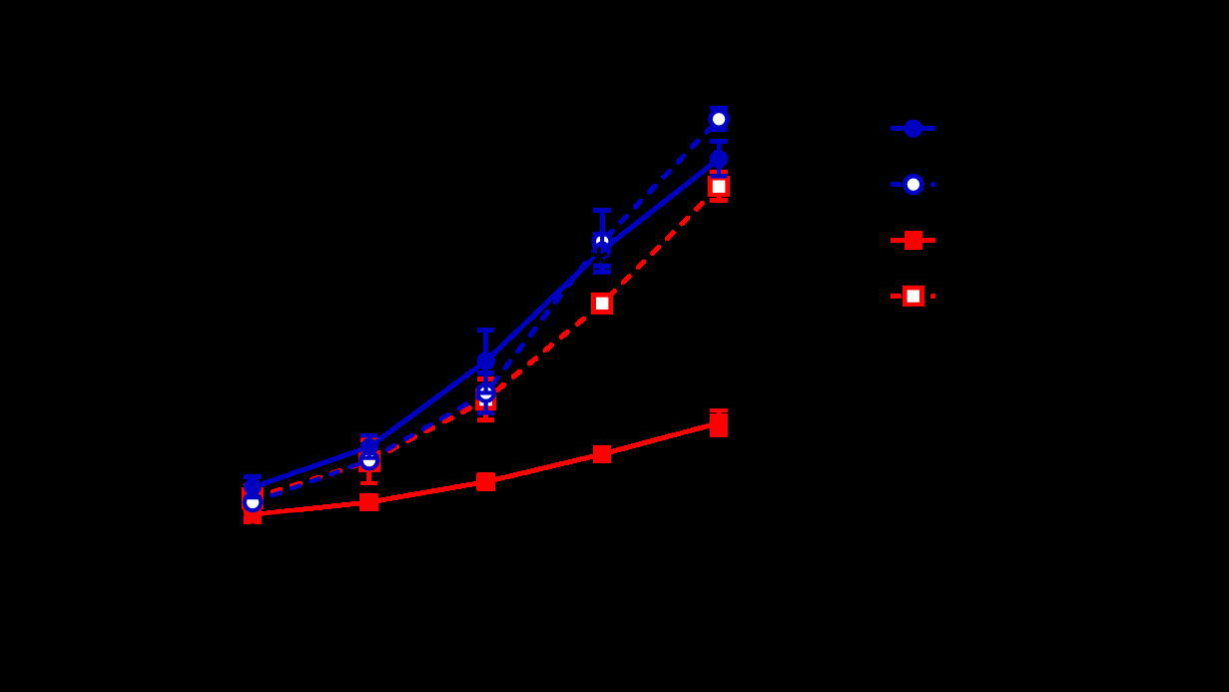
Macrophage depletion restores sensory vasodilation in mesenteric arteries with IBD. Data are mean ± SE % maximum dilations elicited by sensory nerves via electrical field stimulation (1-16 Hz) in arteries from Control (blue) and IBD (red) mice treated with PBS liposomes (solid lines) or clodronate liposomes (dashed lines). N=4-5 per group. * = P<0.05 vs IBD-PBS.

#### 4.4.3 Macrophage depletion does not alter the endothelium-independence of sensory vasodilation

Because sensory vasodilation has been reported to include both endothelial dependent and independent mechanisms, we repeated sensory vasodilation measurements in isolated arteries before (PSS) and after incubation in the nitric oxide synthase inhibitor L-NAME (100 µM) (**Figure 6**). Although significant differences in vasodilation were observed at specific stimulation frequencies within groups, statistical analysis revealed that L-NAME treatment was not a significant source of variation in sensory vasodilation arteries from CONT-PBS (**Figure 6A**), CONT-CLOD (**Figure 6B**), IBD-PBS (**Figure 6C**) or IBD-CLOD (**Figure 6D**) groups.

**Figure 6.**
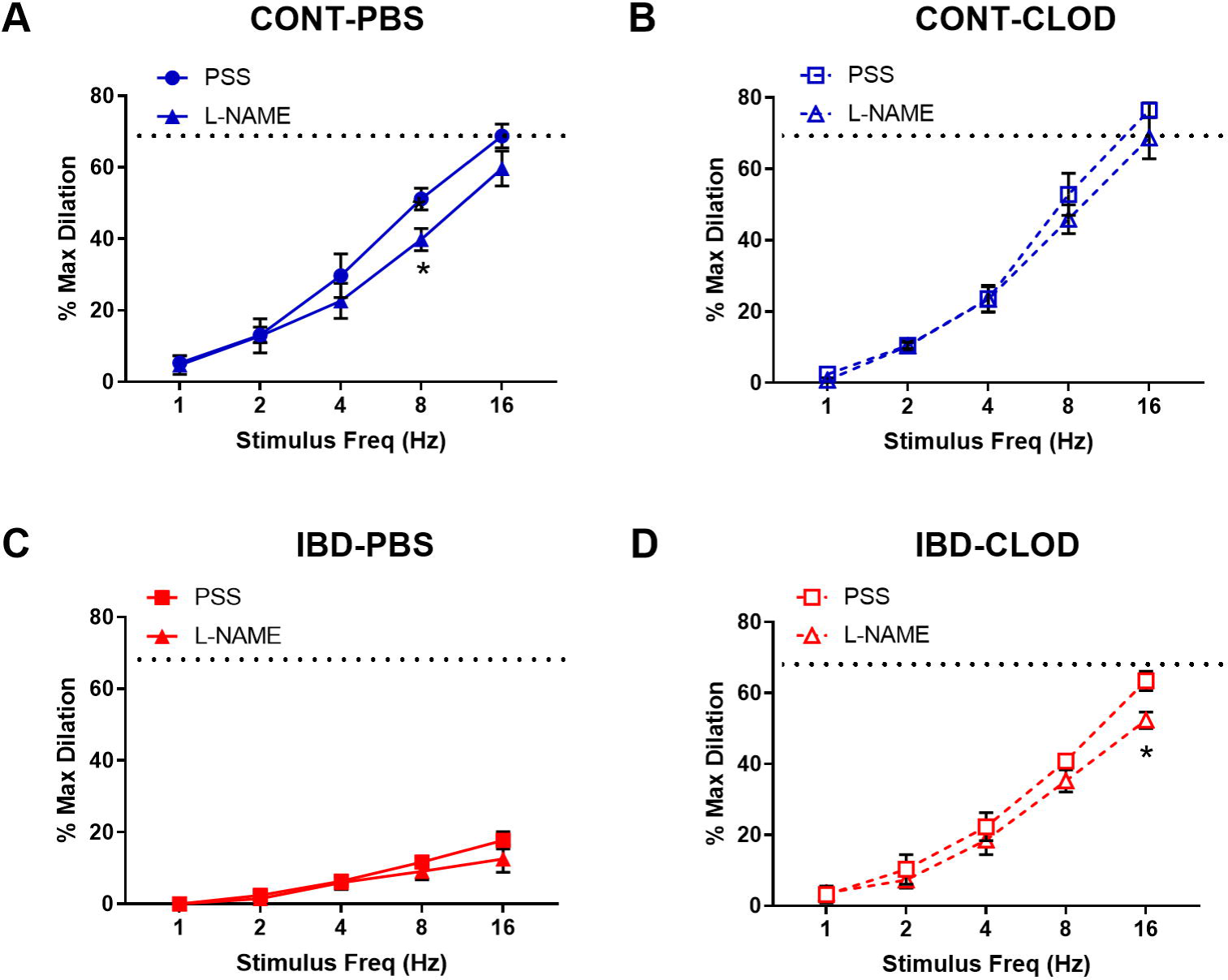
Sensory vasodilation is mostly endothelium-independent and unaffected by macrophage depletion. Data mean ± SE % maximum dilations elicited by sensory nerves via electrical field stimulation (1-16 Hz) in arteries before (PSS) and after incubation in the nitric oxide synthase inhibitor L-NAME (100 µ) from A) CONT-PBS: Control mice treated with PBS liposomes, B) CONT-CLOD: Control mice treated with clodronate liposomes, C) IBD-PBS: IBD mice treated with PBS liposomes, and D) IBD-CLOD: IBD mice treated with clodronate liposomes. Dashed line indicates the max sensory vasodilation reached in CONT-PBS arteries in PSS, which was significantly greater (P<0.05) than max dilation in IBD-PBS but not CONT-PBS or IBD-CLOD. N=4-5 per group. * = P<0.05 vs equivalent stimulation in PSS.

### 4.5 Macrophage depletion improves endothelium-mediated vasodilation in mesenteric arteries with IBD

Because our previous studies found that IBD desensitized endothelium-dependent vasodilation of mesenteric arteries to acetylcholine (ACh) (17), we tested whether macrophage depletion affected ACh-mediated vasodilation. Consistent with the previous study, mesenteric arteries from IBD-PBS mice displayed a significant decrease in maximum ACh dilation and sensitivity (mean EC50s 10^-7.1^ µM vs 10^-6.9^ µM, respectively) compared to CONT-PBS mice (**Figure 7**). Macrophage depletion did not alter ACh concentration-response curves in arteries from CONT-CLOD vs CONT-PBS mice. In contrast, macrophage depletion increased ACh potency ad sensitivity in IBD-CLOD vs IBD-PBS mice (**Figure 7**).

**Figure 7.**
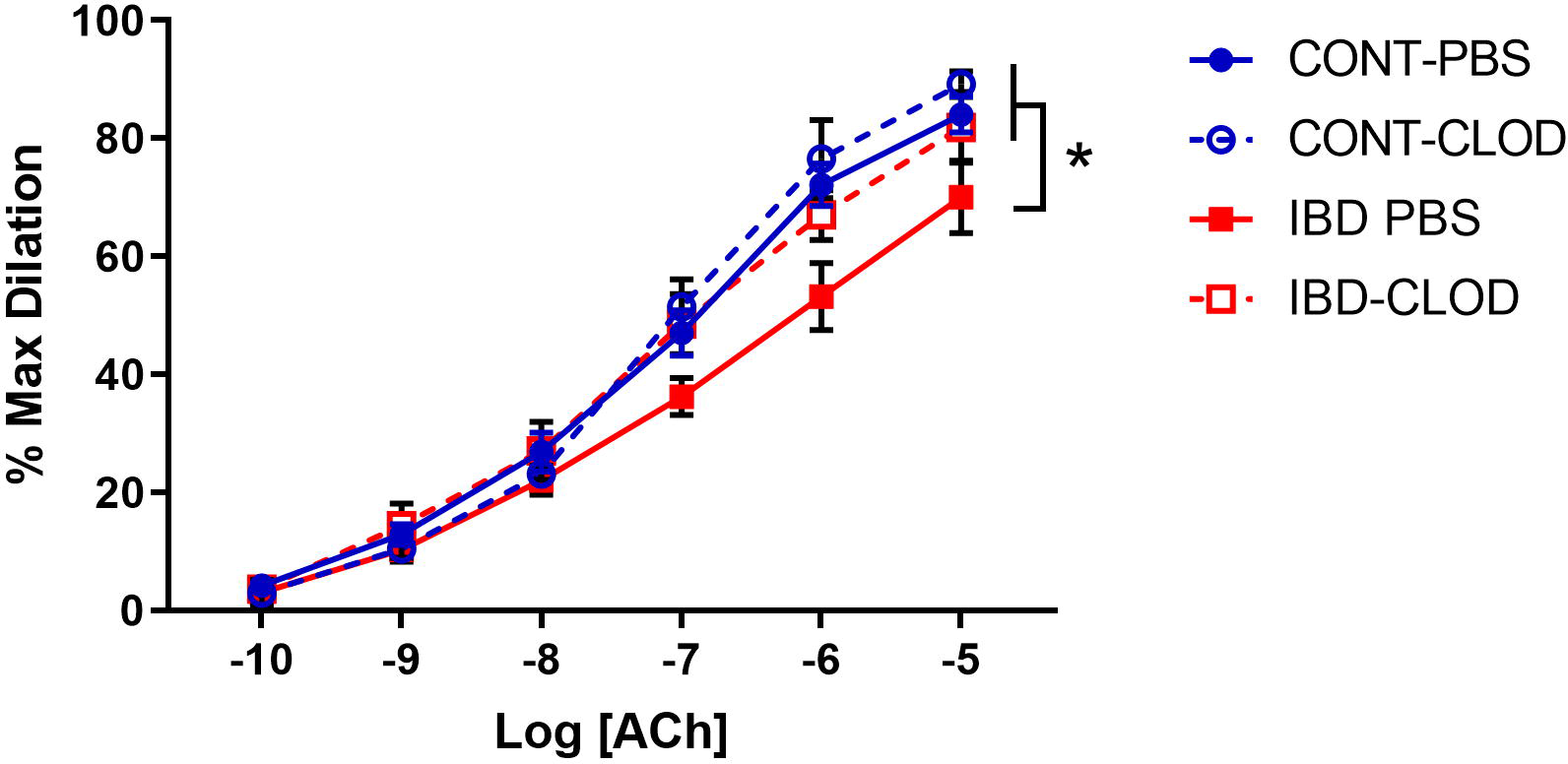
Macrophage depletion improves attenuated ACh dilation in mesenteric arteries with IBD. Data are Data mean ± SE % maximum dilations to increasing concentrations of Acetylcholine (ACh) in preconstricted (phenylephrine, 10µM) mesenteric arteries from Control (blue) and IBD (red) mice treated with PBS liposomes (solid lines) or clodronate liposomes (dashed lines). LogEC50 for each group (in M) are CONT-PBS: -7.08, CONT-CLOD: -7.14, IBD-PBS: -6.90 and IBD-CLOD: -7.20. N=4-5 per group. * = P<0.05 vs IBD-PBS.

### 4.6 Effect of macrophage depletion on perivascular nerve density

Because macrophage depletion was associated with significant improvements in sympathetic vasodilation and sensory vasodilation, we used quantitative immunofluorescence to test whether clodronate treatment altered perivascular nerve density (**Figure 8**). Analysis of max z-projections (**Figure 8A**) revealed no differences in total perivascular nerve density (PGP9.5+) of sympathetic nerve density (tyrosine hydroxylase, TH+) between CONT-PBS, CONT-CLOD, IBD-PBS and IBD-CLOD groups (**Figure 8B,C**). Sensory nerve density, measured with both CGRP+ and SP+ labelling was not different in CONT-PBS vs IBD-PBS arteries (**Figure 8D-E**). Macrophage depletion did not alter CGRP+ or SP+ labeling in CONT-PBS vs CONT-CLOD arteries or IBD-PBS vs IBD-CLOD arteries. However, both CGRP+ and SP+ labeling was significantly increased in IBD-CLOD vs CONT-PBS arteries (**Figure 8D-E**).

**Figure 8.**
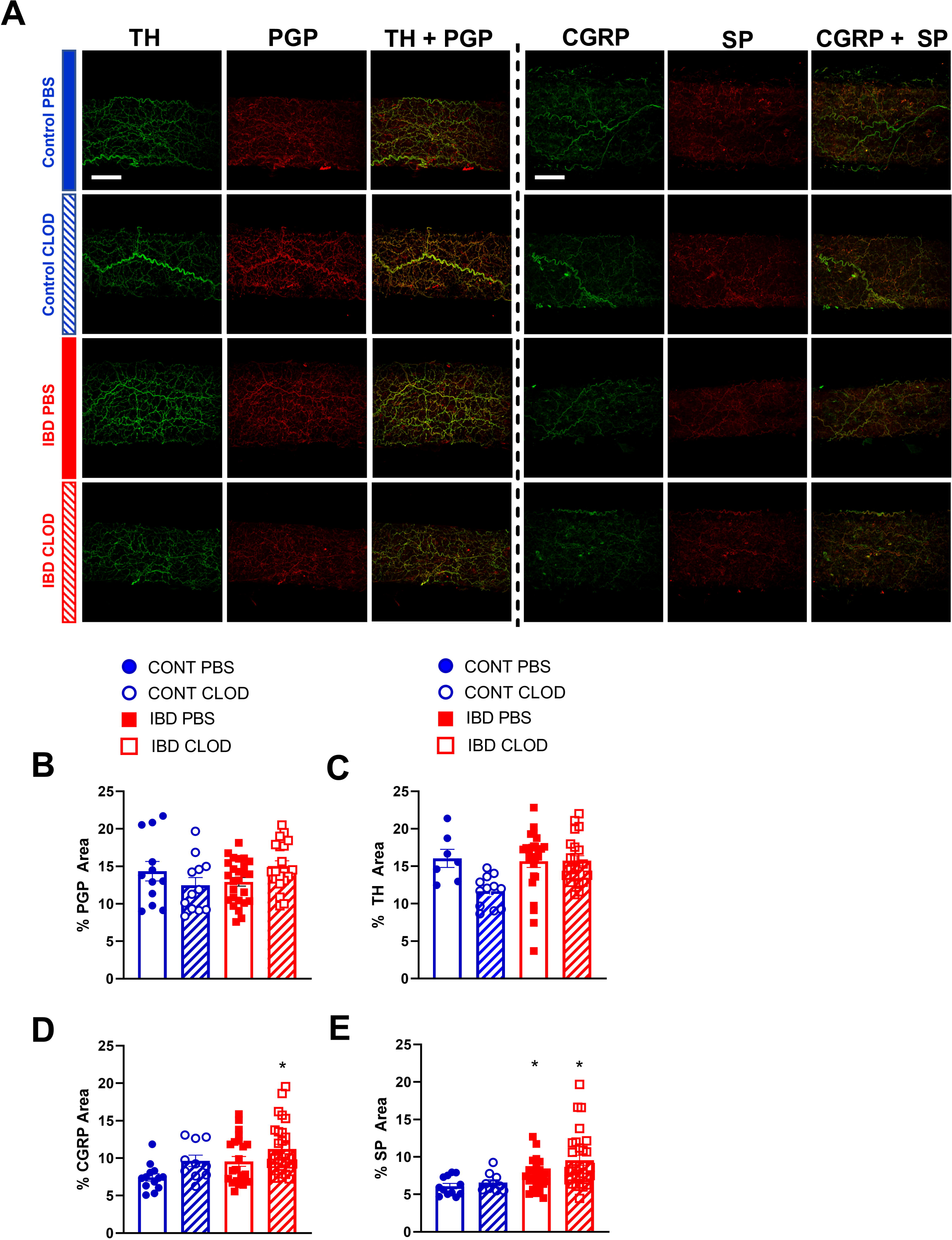
Macrophages depletion increases fluorescence area of sensory neurotransmitters but not the total perivascular nerves or perivascular sympathetic nerves. A) representative max z-projections of perivascular nerves labels from two sets of immunolabeling: perivascular sympathetic nerves (tyrosine hydroxylase, TH, green) + all perivascular nerves (PGP 9.5, red) and sensory neurotransmitters calcitonin gene related peptide (CGRP, green) + substance P (SP, red). Both sets are shown in (top-to-bottom) Control-PBS, Control-CLOD, IBD-PBS and IBD-CLOD groups. Scale bar = 100µm). B-E) quantitation of nerve density. Data are mean ± SE fluorescence area B) all perivascular nerves (PGP 9.5), C) perivascular sympathetic nerves (TH), and D-E) sensory neurotransmitters CGRP and SP of confocal z-projections through the mesenteric artery adventitia of vessels from Control (blue, circles markers) and IBD (red, squares) mice treated with PBS (open bars), or clodronate (striped bars) liposomes. N = 10-25 images from 3-4 mice per group.* = P<0.05 vs CONT-PBS.

### 4.7. Sensory neurotransmitter expression in perivascular macrophages increases with IBD

Macrophage accumulation with IBD impaired, and macrophage depletion reversed perivascular nerve dysfunction in mesenteric arteries with IBD, thus we used confocal immunolabeling and colocalization analysis to determine whether IBD was associated with an increase in the association of the macrophage protein F4/80 and the sensory neurotransmitter receptor proteins RAMP1 and NK1R (**Figure 9A**). Colocalization analysis showed that in the mesenteric artery adventitia, IBD was associated with increased colocalization area of macrophages (F4/80) with sensory neurotransmitter receptor proteins (RAMP1 and NK1, **Figure 9B**). Because nuclear labeling was evident (**Figure 9A**), we also measured colocalization within the nuclei and found IBD-related increased in nuclear colocalization (**Figure 9C**). In addition to the area-based colocalization measurements, fluorescence intensity of colocalized pixels was measured, showing increased colocalized intensity in both whole macrophages (Figure 9D) and macrophage nuclei (**Figure 9D**).

**Figure 9.**
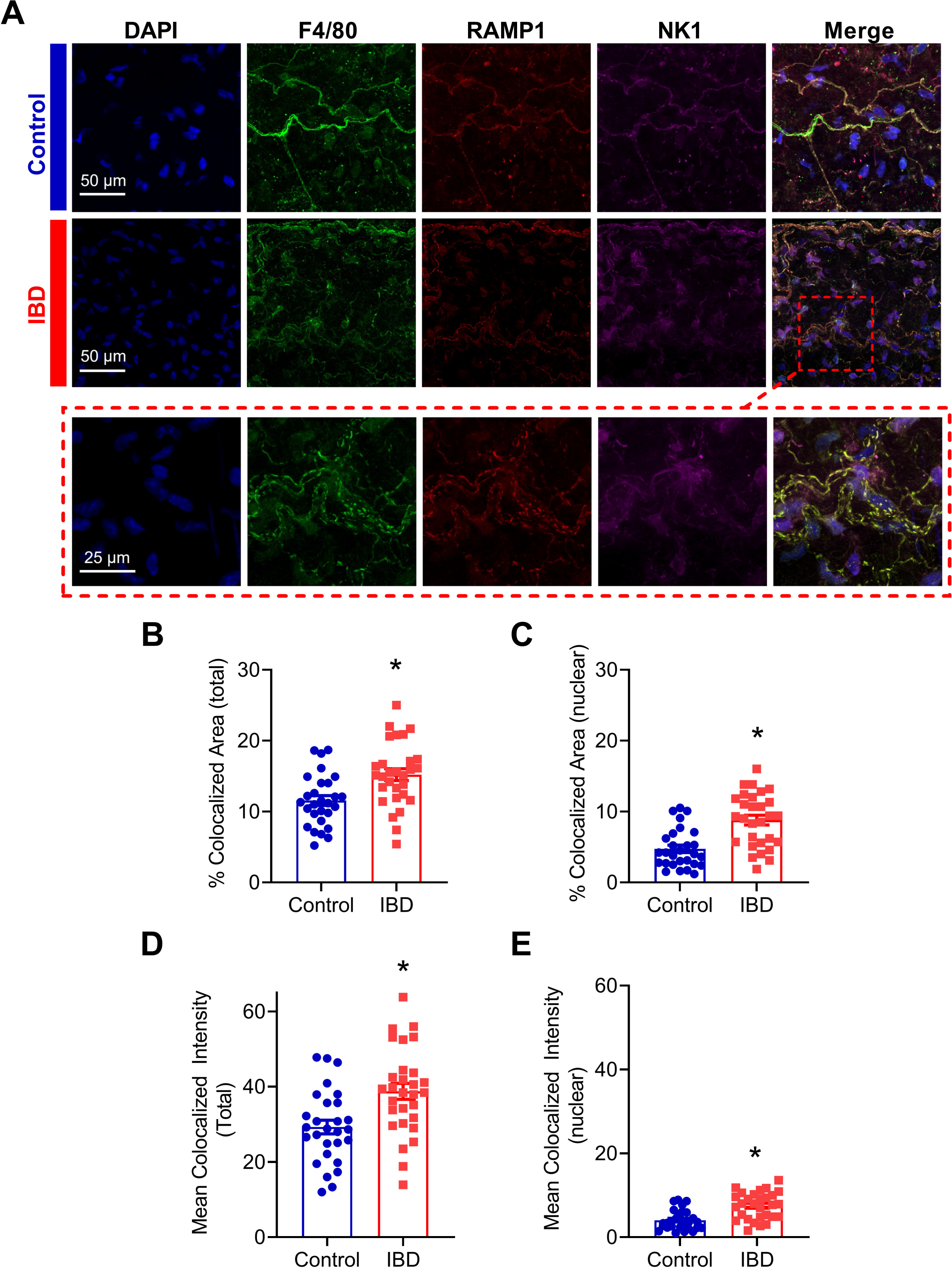
Adventitial macrophages express more sensory neurotransmitter receptors with IBD. A) Representative max z-projections of the vascular adventitia of mesenteric arteries Control (Top, blue label) and IBD (Center, red label) and a zoomed inset from IBD (Bottom, dashed red box) labeled for nuclei, macrophages (F4/80, green), CGRP receptors (RAMP1) and SP receptors (NK1R). B-C) Area of colocalization of RAMP1 and NK1R with (B) total macrophage area and (C) nuclear region of macrophages. D-E) Fluorescence intensity of colocalized pixels in (D) total macrophage area (E) and macrophage nuclei. N = 28-29 images from 3-4 mice per group. * = P<0.05 vs Control.

## 5. Discussion

### 5.1 Summary

The present study used an established IL10^-\-^ + *H. hepaticus* mouse model of IBD (31) to define the mechanisms by which perivascular nerve function in mesenteric arteries is significantly impaired with IBD, likely contributing to the restricted intestinal blood flow often observed in patients. RNA sequencing and analysis of differentially expressed genes from IBD vs Control arteries identified adventitial leukocytes and, more specifically, macrophages as a potential driver of perivascular nerve dysfunction in mesenteric arteries with IBD. Quantitative analysis of confocally-imaged immunolabeled arteries showed that leukocytes, including a significant population of CD68+ and CD163+ macrophages, accumulate in the arterial adventitia near perivascular nerves with IBD. Further, this macrophage population was successfully eliminated following intraperitoneal injection of clodronate liposomes. Next, clodronate or PBS liposome treatment was utilized to determine the effect of macrophage depletion in perivascular nerve function in isolated mesenteric arteries. Using pressure myography and electrical field stimulation to define sympathetic and sensory nerve function, our key findings are that elimination of adventitial macrophages was associated with restoration of (1) sensory vasodilation, (2) sensory inhibition of sympathetic vasoconstriction, and (3) sympathetic vasoconstriction. Sensory vasodilation in this model and its improvements following macrophage depletion appear to be mostly endothelium independent, and they also occur in the absence of major changes in nerve density. Based on the current data, we suggest that impaired sensory vasodilation in mesenteric arteries during IBD occurs due to aberrant adventitial neuro-immune signaling between perivascular nerves and the nearby expanded macrophage population.

### 5.2 Macrophage accumulation in the vascular adventitia with IBD

Adventitial macrophage accumulation and/or polarization has been described in both conduit and resistance arteries from a wide variety of diseases, including autoimmune diseases (see review (32)), but this is the first study to measure the adventitial macrophage population with IBD or investigate its effect on perivascular nerve function in IBD. RNA sequencing of intact mesenteric arteries identified multiple macrophage markers and macrophage-associated genes among the genes with the most increased expression with IBD. Examination of GO terms associated with differentially expressed genes also pointed to leukocytes, macrophages and associated inflammation as a potential target in vascular dysfunction with IBD (**Figure 2, Supplemental Table 1**). Based on this data, and because macrophages have been identified as the most abundant cells in the adventitia in health and disease (33), we pursued macrophage infiltration as a potential cause of perivascular nerve dysfunction. In RNA sequencing, IBD-upregulated genes included pan-macrophage markers, genes associated with both classical (M1-like) and alternative activation (M2-like), and genes associated with macrophage chemotaxis. Upregulated pan macrophage markers included *adgre1, itgam* and *csf1r*, which have each been identified as putative macrophage markers in the vascular adventitia (33). Upregulated genes associated with an M1-like phenotype included *ccl9*, *ccl5* and *cxcl9* (34). Immunolabeling for MHCII and CD68 further indicated an increase in total and M1-like macrophages (**Figure 3**). Upregulated genes associated with M2-like phenotype included *cd163*, *mrc1, ms4a4a* (35), with immunolabeling for CD163 also increased in IBD. Collectively, these data support a broad expansion of arterial macrophages of multiple phenotypes in IBD. Although a more nuanced characterization of adventitial macrophage phenotype is relevant to both IBD and vascular inflammation, the major purpose of the present study was to functionally define the relationship between the overall adventitial macrophage population and perivascular nerve function in IBD. An important focus for ongoing work will be to determine whether macrophage expansion in the adventitia with IBD represents expansion of the resident population or recruitment of new monocyte-derived macrophages, as both populations are present in the healthy adventitia (33). In support of expanded resident macrophage population, we found that many genes with increased expression in arteries with IBD (*ms4a4a, ms4a6c, il1r1, cd36*, **Figure 3**, **Supplemental Table 1**) overlap with genes enriched in peripheral nervous system (PNS) macrophages, (36), which are self-renewing. The overlap between the gene signatures of IBD adventitia and PNS macrophages is also interesting because both macrophage populations reside in close proximity to a nerve microenvironment. Supporting an expanded population of bone marrow derived macrophages, we found increased expression of *s100a9* in mesenteric arteries, which is associated with early monocyte to macrophage differentiation (33).

### 5.3 Adventitial nerve-macrophage signaling in IBD

Consistent with previous work in our laboratory (17), this study found that IBD was associated with drastically impaired sensory vasodilation (**Figure 5**) and impaired inhibition of sympathetic vasoconstriction, despite depressed sympathetic vasoconstriction in IBD (**Figure 4A,C**). Surprisingly, all three vasomotor functions were completely restored following macrophage depletion with clodronate liposomes (**Figures 4-5**). These data highlight that the net result of adventitial macrophage signaling across all phenotypes is impaired perivascular nerve function.

Most work on neuro-immune interactions within and outside the vasculature has focused on sympathetic nerves, and the positive correlation between sympathetic outflow and inflammation throughout the body is well-defined in many diseases. In IBD, studies in human intestine have demonstrated a correlation between increased MHCII+ immune cell presence and decreased enteric nerves function (37) and density (38). Counterintuitively, our study found that IBD was associated with increased adventitial macrophages (**Figure 3**) but decreased sympathetic vasoconstriction, which was restored following macrophage depletion (**Figure 4**); sympathetic nerve density remained unchanged (**Figure 8**). Few studies have investigated the macrophage-nerve relationship in the mesenteric vasculature. Studies in a rat model of hypertension found that CD163+ macrophage accumulation in the mesenteric artery adventitia is linked to increased blood pressure through macrophage-derived oxidate stress acting on ganglionic neuronal alpha-2 adrenergic receptors (39). In that study, clodronate-based macrophage depletion acted to lower blood pressure by preserving neuronal alpha-2 receptor function. Here, increased CD163+ macrophage population in the mesenteric artery adventitia (**Figures 3**) is associated with decreased sympathetic nerve function, suggesting that macrophages may mediate vascular dysfunction in distinct ways during IBD versus hypertension and other cardiovascular diseases.

Macrophage accumulation appears to have a much greater and more detrimental effect on perivascular sensory nerves compared to sympathetic nerves. In earlier studies(17), we found that despite impaired sensory vasodilation with IBD, there was not a significant decrease in the content or release of the sensory neurotransmitters CGRP and substance P. Similarly, the function of CGRP receptors, the primary vasodilatory peptide, released was unaltered with IBD, as shown by exogenous CGRP concentration-response curves. SP vasodilation was impaired despite an increase in NK1 receptor expression with IBD. Norepinephrine sensitivity was also unchanged with IBD (17). Considering the seemingly preserved neurotransmission and largely preserved receptor function with unchanged nerve density (**Figure 8**), what is happening to the released neurotransmitter? We propose that rather than acting downstream to affect vasodilation, released CGRP and substance P binds to receptors expressed on the expanded population of closely-approximated macrophages. Thus, sensory transmitter released from nerves would mediate immune-mediated effects in place of the vasodilation observed in arteries with a healthy adventitial macrophage population. Consistent with this hypothesis, we found that in addition to an increased population of adventitial macrophages, IBD was associated with increased expression of CGRP and SP receptors within those adventitial macrophages (**Figure 9**). Confocal images also highlighted that macrophage projections appear to closely approximate perivascular nerve structures, with CGRP and SP receptors expressed in these approximations (**Figure 9A**). The possibility that macrophages can effectively “steal” released neurotransmitters to affect inflammatory processes is supported by multiple areas of research. Both CGRP and substance P can affect macrophage phenotype and the release of inflammatory mediators in vitro (40, 41, 42), although the effects of each neuropeptide on perivascular macrophages is unknown. Moreover, local macrophage activation by neuronally released neurotransmitter was recently defined in the intestinal wall. In a series of experiments, Gabanyi et al (43) characterized a population of intestinal macrophages expressing β2 adrenergic receptors that are uniquely located near nerves and activated by norepinephrine released from extrinsic nerves to induce M2 polarization. This concept is consistent with our findings of depressed sympathetic vasoconstriction (**Figure 4**) and increased gene and protein indicators of M2 polarization (**Figures 2-3**) with IBD. The gene expression profiling from the Gabanyi study further found that the CGRP receptor protein RAMP1 was the only other neurotransmitter receptor with increased expression in the muscularis macrophage population, although follow-up imaging studies were not performed. Given that perivascular nerves on mesenteric arteries extend into the intestinal wall (44), they may similarly affect the local macrophage population that express neurotransmitter receptors.

### 5.4 Role of endothelium and nitric oxide in perivascular nerve dysfunction

The role of the endothelium in both sensory vasodilation and IBD pathogenesis is unclear and somewhat controversial. In both health and disease, sensory vasodilation has been described as mostly endothelium dependent (45, 46), partially endothelium dependent (47, 48) and endothelium independent (49, 50). Previous work in mouse mesenteric arteries supports partial endothelial dependence as well as increased endothelial dependence with aging (48), suggesting that the mechanism of sensory vasodilation in mesenteric arteries can change based on pathology. In the present study, we examined changes in nitric oxide (NO)-dependent mechanisms of sensory vasodilation, as NO-mediated endothelial dysfunction is evident within the intestinal wall (15). We found that blocking nitric oxide synthase with L-NAME had a negligible effect on sensory vasodilation in vessels from Control and IBD mice with intact macrophages (**Figure 6 A,C**). L-NAME also had no effect on sensory vasodilation in Control or IBD mice after macrophage depletion (**Figure 6 B,D**), suggesting that nitric oxide generation is not an important target for restoring sensory vasodilation in this model. Because CGRP and SP can also elicit vasodilation via endothelium dependent hyperpolarization in these arteries (51), additional experiments exploring these mechanisms would be required to fully define how IBD affects endothelial effects of CGRP and SP.

Independent of sensory nerve function, endothelial dysfunction of varying severity and mechanism has been demonstrated in murine (52) and human (53) studies of IBD. Here, we found a small but significant decrease in acetylcholine-mediated dilation in mesenteric arteries with IBD (**Figure 7**). Macrophage depletion restored acetylcholine responses in arteries from IBD mice, suggesting that macrophage-mediated endothelial dysfunction contributes to vascular dysfunction in IBD independent of perivascular nerve function. The presence of some endothelial dysfunction in this model is consistent with the IBD-related increase in expression of the chemokine *ccl5* and its receptor *ccr1*, which are linked to endothelial dysfunction and leukocyte transmigration into arteries (54). Collectively, our data suggest that “inside-out” vascular dysfunction arising from the endothelium is less significant in IBD than “outside-in” mechanisms initiated within the adventitia.

### 5.5 Limitations

This study clearly demonstrates the presence of significant macrophage-associated perivascular nerve dysfunction, but there are also limitations to the study that warrant further discussion. It is important to note that no mouse model of IBD can currently recapitulate all aspects of human IBD. The IL10^-/-^ model is widely used and accepted (55) but has limiting factors. The use of the IL10^-/-^ model may affect the immune responses observed, as immune cells cannot produce and respond to IL10 in the same way as human IBD patients. In addition, the severity of IBD in IL10^-/-^ is quite variable across studies, and as disease severity varies with microbiome differences arising from mouse strain, vendor and specific housing conditions (56). Clodronate studies also have limitations. The use of clodronate to deplete macrophages may have indeterminate effects on vascular function independent of the elimination of adventitial macrophages, as it depletes macrophages throughout the body. We also halved doses 2 and 3 of clodronate, as repeating the original dose led to ∼25% mortality in the control group (consistent with published studies (57, 58)) and up to 50% mortality in the IBD group. No studies using clodronate in IL10^-/-^ mice have reported mortality. Macrophage phenotyping is also a limitation in this study, as the leukocyte phenotyping used for immunofluorescence was relatively rudimentary. While it effectively shows accumulations of very broad populations of cells, it remains a non-definitive classification of macrophage subtypes. Follow-up studies with flow cytometry and/or single cell sequencing will be needed to more rigorously define the adventitial immune cell population with IBD.

### 5.6 Conclusions

Collectively, our findings show for the first time that expansion of the adventitial macrophage population adversely impacts the function of perivascular nerves in inflammatory bowel disease, where impaired blood flow to the intestine is common. Sensory vasodilation, which is dramatically impaired in mesenteric arteries with IBD can be restored following macrophage depletion. Moreover, the IBD-expanded population of adventitial macrophages has increased expression of receptors for the sensory neurotransmitters CGRP and substance P. We propose that during IBD, CGRP and SP released from sensory nerves is functionally sequestered via macrophage binding, leading to decreased sensory vasodilation and increased macrophage activation. Overall, targeting perivascular macrophages may represent a promising target to promote improved mesenteric blood flow with inflammatory bowel disease.

## Data availability statement

The original contributions presented in the study are included in the article/supplementary material, and further inquiries can be directed to the corresponding author. All RNA sequencing data has been made publicly available at https://doi.org/10.7910/DVN/JEQ6ND.

## Author contributions

EB conceived and designed the experiments. EB, EG, OL and BJ performed the experiments. EB, BJ, OL TJ, KW and SS analyzed the data. All authors approved the final version of the manuscript.

