## Supplemental Table for "Adventitial macrophage accumulation impairs perivascular nerve function in mesenteric arteries with inflammatory bowel disease"

**Supplementary Table 1. Complete listing of differentially expressed genes in mesenteric arteries from IBD vs Control mice.** Data show each of the differentially expressed genes listed from highest to lowest fold change in expression from IBD vs Control arteries with associated P and Q values. All genes shown have Q values < 0.05.

| Gene Name | Target | Log2 Fold Change | Fold Change | P value | Q value |
| --- | --- | --- | --- | --- | --- |
| Jchain | immunoglobulin joining chain | 3.67133 | **12.74034** | 0.00005 | 0.00621 |
| Ms4a4a | membrane-spanning 4-domains, subfamily A, member 4A | 2.90959 | **7.51403** | 0.00005 | 0.00621 |
| Cd163 | CD163 antigen | 2.35215 | **5.10583** | 0.00005 | 0.00621 |
| Mt2 | metallothionein 2 | 2.28306 | **4.86708** | 0.00005 | 0.00621 |
| Add2 | adducin 2 (beta) | 2.27127 | **4.82748** | 0.00005 | 0.00621 |
| Pcdh20 | protocadherin 20 | 2.24675 | **4.74611** | 0.00005 | 0.00621 |
| Ccl9 | chemokine (C-C motif) ligand 9 | 2.20597 | **4.61385** | 0.00005 | 0.00621 |
| Ms4a6c | membrane-spanning 4-domains, subfamily A, member 6C | 2.15601 | **4.45681** | 0.00005 | 0.00621 |
| Fkbp5 | FK506 binding protein 5 | 2.01816 | **4.05066** | 0.00005 | 0.00621 |
| Siglec1 | sialic acid binding Ig-like lectin 1, sialoadhesin | 2.00184 | **4.00511** | 0.00005 | 0.00621 |
| Ms4a6d | membrane-spanning 4-domains, subfamily A, member 6D | 1.92841 | **3.80635** | 0.00005 | 0.00621 |
| Hif3a | hypoxia inducible factor 3, alpha subunit | 1.78961 | **3.45722** | 0.00005 | 0.00621 |
| Ccr1 | chemokine (C-C motif) receptor 1 | 1.78147 | **3.43776** | 0.00005 | 0.00621 |
| Itgam | integrin alpha M | 1.71530 | **3.28365** | 0.00005 | 0.00621 |
| C5ar1 | complement component 5a receptor 1 | 1.65530 | **3.14989** | 0.00005 | 0.00621 |
| Cd33 | CD33 antigen | 1.64062 | **3.11799** | 0.00005 | 0.00621 |
| Tlr7 | toll-like receptor 7 | 1.62461 | **3.08358** | 0.00005 | 0.00621 |
| Ifi205 | interferon activated gene 205 | 1.60962 | **3.05172** | 0.00005 | 0.00621 |
| Igkc | immunoglobulin kappa constant | 1.60187 | **3.03536** | 0.00005 | 0.00621 |
| Adgre1 | adhesion G protein-coupled receptor E1 | 1.59667 | **3.02444** | 0.00005 | 0.00621 |
| Cd300a | CD300A molecule | 1.59253 | **3.01577** | 0.00015 | 0.01595 |
| Ccl6 | chemokine (C-C motif) ligand 6 | 1.58953 | **3.00951** | 0.00005 | 0.00621 |
| Galnt15 | polypeptide N-acetylgalactosaminyltransferase 15 | 1.58463 | **2.99931** | 0.00005 | 0.00621 |
| Pla2g7 | phospholipase A2, group VII (platelet-activating factor acetylhydrolase, plasma) | 1.57960 | **2.98887** | 0.00005 | 0.00621 |
| Timd4 | T cell immunoglobulin and mucin domain containing 4 | 1.57629 | **2.98201** | 0.00010 | 0.01142 |
| Wdfy4 | WD repeat and FYVE domain containing 4 | 1.54548 | **2.91901** | 0.00005 | 0.00621 |
| F13a1 | coagulation factor XIII, A1 subunit | 1.54299 | **2.91398** | 0.00005 | 0.00621 |
| Csf1r | colony stimulating factor 1 receptor | 1.53024 | **2.88834** | 0.00005 | 0.00621 |
| Clec10a | C-type lectin domain family 10, member A | 1.50107 | **2.83052** | 0.00060 | 0.04848 |
| Trim10 | tripartite motif-containing 10 | 1.48733 | **2.80370** | 0.00005 | 0.00621 |
| Coro2a | coronin, actin binding protein 2A | 1.47983 | **2.78916** | 0.00015 | 0.01595 |
| March1 | membrane-associated ring finger (C3HC4) 1 | 1.47507 | **2.77997** | 0.00010 | 0.01142 |
| Usp54 | ubiquitin specific peptidase 54 | 1.47156 | **2.77322** | 0.00005 | 0.00621 |
| Mrc1 | mannose receptor, C type 1 | 1.45170 | **2.73530** | 0.00005 | 0.00621 |
| Zbtb16 | zinc finger and BTB domain containing 16 | 1.43610 | **2.70588** | 0.00005 | 0.00621 |
| Fcgr3 | Fc receptor, IgG, low affinity III | 1.43568 | **2.70510** | 0.00005 | 0.00621 |
| 9330159F19Rik | RIKEN cDNA 9330159F19 gene | 1.43230 | **2.69876** | 0.00005 | 0.00621 |
| Mt1 | metallothionein 1 | 1.39260 | **2.62551** | 0.00005 | 0.00621 |
| Tsc22d3 | TSC22 domain family, member 3 | 1.39126 | **2.62307** | 0.00005 | 0.00621 |
| Mgl2 | macrophage galactose N-acetyl-galactosamine specific lectin 2 | 1.38832 | **2.61774** | 0.00005 | 0.00621 |
| C3ar1 | complement component 3a receptor 1 | 1.36775 | **2.58068** | 0.00020 | 0.02011 |
| Per2 | period circadian clock 2 | 1.34267 | **2.53621** | 0.00005 | 0.00621 |
| Gm8812 | predicted gene 8812 | 1.33763 | **2.52736** | 0.00025 | 0.02474 |
| Fam13a | family with sequence similarity 13, member A | 1.32786 | **2.51031** | 0.00005 | 0.00621 |
| Fmo3 | flavin containing monooxygenase 3 | 1.31790 | **2.49303** | 0.00005 | 0.00621 |
| Lvrn | laeverin | 1.29559 | **2.45477** | 0.00005 | 0.00621 |
| Il1rl1 | interleukin 1 receptor-like 1 | 1.28984 | **2.44501** | 0.00035 | 0.03159 |
| Adamts14 | a disintegrin-like and metallopeptidase (reprolysin type) with thrombospondin type 1 motif, 14 | 1.28763 | **2.44126** | 0.00005 | 0.00621 |
| Lrg1 | leucine-rich alpha-2-glycoprotein 1 | 1.28012 | **2.42860** | 0.00005 | 0.00621 |
| Pdk4 | pyruvate dehydrogenase kinase, isoenzyme 4 | 1.27521 | **2.42034** | 0.00005 | 0.00621 |
| Slc10a6 | solute carrier family 10 (sodium/bile acid cotransporter family), member 6 | 1.25997 | **2.39490** | 0.00005 | 0.00621 |
| Rasd1 | RAS, dexamethasone-induced 1 | 1.24155 | **2.36453** | 0.00005 | 0.00621 |
| Car2 | carbonic anhydrase 2 | 1.23834 | **2.35927** | 0.00045 | 0.03819 |
| Aqp1 | aquaporin 1 | 1.21680 | **2.32431** | 0.00005 | 0.00621 |
| C1qc | complement component 1, q subcomponent, C chain | 1.21368 | **2.31928** | 0.00005 | 0.00621 |
| Mia3 | melanoma inhibitory activity 3 | 1.19697 | **2.29257** | 0.00005 | 0.00621 |
| Rergl | RERG/RAS-like | 1.19455 | **2.28873** | 0.00005 | 0.00621 |
| Clec4a1 | C-type lectin domain family 4, member a1 | 1.19328 | **2.28672** | 0.00025 | 0.02474 |
| Gypa | glycophorin A | 1.18831 | **2.27885** | 0.00030 | 0.02818 |
| Clca3a1 | chloride channel accessory 3A1 | 1.17285 | **2.25457** | 0.00005 | 0.00621 |
| Sult1a1 | sulfotransferase family 1A, phenol-preferring, member 1 | 1.16602 | **2.24391** | 0.00005 | 0.00621 |
| Glns-ps1 | glutamine synthetase pseudogene 1 | 1.16260 | **2.23860** | 0.00005 | 0.00621 |
| C1qa | complement component 1, q subcomponent, alpha polypeptide | 1.15587 | **2.22818** | 0.00005 | 0.00621 |
| Cyp2e1 | cytochrome P450, family 2, subfamily e, polypeptide 1 | 1.13743 | **2.19988** | 0.00005 | 0.00621 |
| Fam107a | family with sequence similarity 107, member A | 1.12338 | **2.17856** | 0.00005 | 0.00621 |
| Glul | glutamate-ammonia ligase (glutamine synthetase) | 1.11632 | **2.16794** | 0.00005 | 0.00621 |
| Ccbe1 | collagen and calcium binding EGF domains 1 | 1.11621 | **2.16777** | 0.00005 | 0.00621 |
| Ctse | cathepsin E | 1.11488 | **2.16577** | 0.00015 | 0.01595 |
| C1qb | complement component 1, q subcomponent, beta polypeptide | 1.11152 | **2.16073** | 0.00005 | 0.00621 |
| Nckap1l | NCK associated protein 1 like | 1.10701 | **2.15399** | 0.00005 | 0.00621 |
| Kcna1 | potassium voltage-gated channel, shaker-related subfamily, member 1 | 1.10008 | **2.14367** | 0.00005 | 0.00621 |
| Lyz2 | lysozyme 2 | 1.09464 | **2.13559** | 0.00005 | 0.00621 |
| Cybb | cytochrome b-245, beta polypeptide | 1.07344 | **2.10444** | 0.00055 | 0.04523 |
| Acss3 | acyl-CoA synthetase short-chain family member 3 | 1.07145 | **2.10154** | 0.00020 | 0.02011 |
| Ifi207 | interferon activated gene 207 | 1.06402 | **2.09076** | 0.00005 | 0.00621 |
| Zfp871 | zinc finger protein 871 | 1.06311 | **2.08943** | 0.00005 | 0.00621 |
| C7 | complement component 7 | 1.06204 | **2.08788** | 0.00005 | 0.00621 |
| Fcrls | Fc receptor-like S, scavenger receptor | 1.05591 | **2.07903** | 0.00015 | 0.01595 |
| Klf15 | Kruppel-like factor 15 | 1.05002 | **2.07056** | 0.00005 | 0.00621 |
| S100a9 | S100 calcium binding protein A9 (calgranulin B) | 1.04322 | **2.06082** | 0.00005 | 0.00621 |
| Ctss | cathepsin S | 1.03248 | **2.04554** | 0.00005 | 0.00621 |
| 9430015G10Rik | RIKEN cDNA 9430015G10 gene | 1.02812 | **2.03937** | 0.00040 | 0.03491 |
| Serpina3n | serine (or cysteine) peptidase inhibitor, clade A, member 3N | 1.02625 | **2.03673** | 0.00005 | 0.00621 |
| Sgms1 | sphingomyelin synthase 1 | 1.02341 | **2.03272** | 0.00005 | 0.00621 |
| S100a8 | S100 calcium binding protein A8 (calgranulin A) | 1.00539 | **2.00748** | 0.00050 | 0.04149 |
| Doc2b | double C2, beta | 0.95849 | **1.94327** | 0.00005 | 0.00621 |
| Pld4 | phospholipase D family, member 4 | 0.95347 | **1.93652** | 0.00005 | 0.00621 |
| Ctsc | cathepsin C | 0.94315 | **1.92272** | 0.00005 | 0.00621 |
| Ptafr | platelet-activating factor receptor | 0.93682 | **1.91431** | 0.00040 | 0.03491 |
| Ccr2 | chemokine (C-C motif) receptor 2 | 0.93303 | **1.90929** | 0.00005 | 0.00621 |
| Cd36 | CD36 molecule | 0.93234 | **1.90837** | 0.00005 | 0.00621 |
| Abca9 | ATP-binding cassette, sub-family A (ABC1), member 9 | 0.92949 | **1.90460** | 0.00005 | 0.00621 |
| Slc7a8 | solute carrier family 7 (cationic amino acid transporter, y+ system), member 8 | 0.92902 | **1.90399** | 0.00030 | 0.02818 |
| Iqgap2 | IQ motif containing GTPase activating protein 2 | 0.91478 | **1.88528** | 0.00005 | 0.00621 |
| Gm16439 | predicted pseudogene 16439 | 0.91155 | **1.88107** | 0.00040 | 0.03491 |
| Tcf23 | transcription factor 23 | 0.90248 | **1.86928** | 0.00005 | 0.00621 |
| Mal | myelin and lymphocyte protein, T cell differentiation protein | 0.89407 | **1.85841** | 0.00005 | 0.00621 |
| Fmo2 | flavin containing monooxygenase 2 | 0.88223 | **1.84323** | 0.00035 | 0.03159 |
| Gpnmb | glycoprotein (transmembrane) nmb | 0.87493 | **1.83392** | 0.00005 | 0.00621 |
| Klf9 | Kruppel-like factor 9 | 0.86619 | **1.82284** | 0.00005 | 0.00621 |
| Rin3 | Ras and Rab interactor 3 | 0.86417 | **1.82030** | 0.00005 | 0.00621 |
| Fcgr2b | Fc receptor, IgG, low affinity IIb | 0.84246 | **1.79310** | 0.00005 | 0.00621 |
| Cd53 | CD53 antigen | 0.83988 | **1.78990** | 0.00060 | 0.04848 |
| Hmgcs2 | 3-hydroxy-3-methylglutaryl-Coenzyme A synthase 2 | 0.81575 | **1.76022** | 0.00005 | 0.00621 |
| Xdh | xanthine dehydrogenase | 0.81121 | **1.75469** | 0.00005 | 0.00621 |
| Myrip | myosin VIIA and Rab interacting protein | 0.81087 | **1.75426** | 0.00005 | 0.00621 |
| Lifr | leukemia inhibitory factor receptor | 0.80610 | **1.74848** | 0.00020 | 0.02011 |
| Crispld1 | cysteine-rich secretory protein LCCL domain containing 1 | 0.80145 | **1.74285** | 0.00030 | 0.02818 |
| Syk | spleen tyrosine kinase | 0.79869 | **1.73952** | 0.00005 | 0.00621 |
| Lyve1 | lymphatic vessel endothelial hyaluronan receptor 1 | 0.79750 | **1.73809** | 0.00005 | 0.00621 |
| Ddit4 | DNA-damage-inducible transcript 4 | 0.79540 | **1.73556** | 0.00005 | 0.00621 |
| Arhgef3 | Rho guanine nucleotide exchange factor (GEF) 3 | 0.78783 | **1.72647** | 0.00005 | 0.00621 |
| Fnip2 | folliculin interacting protein 2 | 0.78601 | **1.72429** | 0.00040 | 0.03491 |
| Cdh19 | cadherin 19, type 2 | 0.78492 | **1.72300** | 0.00005 | 0.00621 |
| Rnf125 | ring finger protein 125 | 0.78411 | **1.72203** | 0.00020 | 0.02011 |
| Rasl10b | RAS-like, family 10, member B | 0.76610 | **1.70067** | 0.00020 | 0.02011 |
| Zfp366 | zinc finger protein 366 | 0.74453 | **1.67543** | 0.00005 | 0.00621 |
| Aldh1a1 | aldehyde dehydrogenase family 1, subfamily A1 | 0.74322 | **1.67391** | 0.00020 | 0.02011 |
| Mlxip | MLX interacting protein | 0.74204 | **1.67254** | 0.00035 | 0.03159 |
| Slco2b1 | solute carrier organic anion transporter family, member 2b1 | 0.73412 | **1.66339** | 0.00005 | 0.00621 |
| Pdpr | pyruvate dehydrogenase phosphatase regulatory subunit | 0.73257 | **1.66160** | 0.00005 | 0.00621 |
| Aph1b | aph1 homolog B, gamma secretase subunit | 0.73224 | **1.66122** | 0.00005 | 0.00621 |
| Plat | plasminogen activator, tissue | 0.72898 | **1.65746** | 0.00005 | 0.00621 |
| Prex2 | phosphatidylinositol-3,4,5-trisphosphate-dependent Rac exchange factor 2 | 0.71223 | **1.63833** | 0.00045 | 0.03819 |
| Osmr | oncostatin M receptor | 0.71199 | **1.63806** | 0.00005 | 0.00621 |
| Ogt | O-linked N-acetylglucosamine (GlcNAc) transferase (UDP-N-acetylglucosamine:polypeptide-N-acetylglucosaminyl transferase) | 0.70794 | **1.63347** | 0.00010 | 0.01142 |
| Fam213a | family with sequence similarity 213, member A | 0.70206 | **1.62683** | 0.00040 | 0.03491 |
| Mmp3 | matrix metallopeptidase 3 | 0.69415 | **1.61793** | 0.00005 | 0.00621 |
| Atp2b2 | ATPase, Ca++ transporting, plasma membrane 2 | 0.68888 | **1.61203** | 0.00005 | 0.00621 |
| Aff1 | AF4/FMR2 family, member 1 | 0.68698 | **1.60992** | 0.00005 | 0.00621 |
| Cmip | c-Maf inducing protein | 0.68336 | **1.60587** | 0.00005 | 0.00621 |
| Fam117b | family with sequence similarity 117, member B | 0.68271 | **1.60516** | 0.00005 | 0.00621 |
| Ptprj | protein tyrosine phosphatase, receptor type, J | 0.68158 | **1.60390** | 0.00020 | 0.02011 |
| Peg3 | paternally expressed 3 | 0.67574 | **1.59741** | 0.00005 | 0.00621 |
| Bcl2l1 | BCL2-like 1 | 0.67077 | **1.59192** | 0.00010 | 0.01142 |
| Nr1d2 | nuclear receptor subfamily 1, group D, member 2 | 0.66614 | **1.58682** | 0.00030 | 0.02818 |
| Tmem2 | transmembrane protein 2 | 0.65953 | **1.57957** | 0.00005 | 0.00621 |
| Rnf152 | ring finger protein 152 | 0.64300 | **1.56157** | 0.00060 | 0.04848 |
| Lnpep | leucyl/cystinyl aminopeptidase | 0.63383 | **1.55168** | 0.00015 | 0.01595 |
| Spock2 | sparc/osteonectin, cwcv and kazal-like domains proteoglycan 2 | 0.62239 | **1.53943** | 0.00005 | 0.00621 |
| Mkl2 | MKL/myocardin-like 2 | 0.62079 | **1.53771** | 0.00040 | 0.03491 |
| S1pr1 | sphingosine-1-phosphate receptor 1 | 0.61959 | **1.53644** | 0.00030 | 0.02818 |
| Frmd4b | FERM domain containing 4B | 0.61801 | **1.53476** | 0.00015 | 0.01595 |
| Tbc1d8b | TBC1 domain family, member 8B | 0.61786 | **1.53460** | 0.00035 | 0.03159 |
| Dll4 | delta-like 4 (Drosophila) | 0.60474 | **1.52071** | 0.00010 | 0.01142 |
| Angptl4 | angiopoietin-like 4 | 0.60122 | **1.51699** | 0.00035 | 0.03159 |
| Slc7a2 | solute carrier family 7 (cationic amino acid transporter, y+ system), member 2 | 0.60023 | **1.51596** | 0.00030 | 0.02818 |
| Setbp1 | SET binding protein 1 | 0.58538 | **1.50043** | 0.00015 | 0.01595 |
| Dapk1 | death associated protein kinase 1 | 0.57746 | **1.49222** | 0.00030 | 0.02818 |
| Lyst | lysosomal trafficking regulator | 0.56609 | **1.48051** | 0.00035 | 0.03159 |
| Jade2 | jade family PHD finger 2 | 0.54889 | **1.46296** | 0.00060 | 0.04848 |
| Mtss1l | metastasis suppressor 1-like | -0.55422 | **0.68103** | 0.00050 | 0.04149 |
| Tnfrsf12a | tumor necrosis factor receptor superfamily, member 12a | -0.57570 | **0.67096** | 0.00030 | 0.02818 |
| Ppp1r3c | protein phosphatase 1, regulatory (inhibitor) subunit 3C | -0.57670 | **0.67050** | 0.00025 | 0.02474 |
| Jund | jun D proto-oncogene | -0.61308 | **0.65380** | 0.00050 | 0.04149 |
| Cd248 | CD248 antigen, endosialin | -0.62838 | **0.64690** | 0.00005 | 0.00621 |
| Micall2 | MICAL-like 2 | -0.63282 | **0.64491** | 0.00035 | 0.03159 |
| Rasl11b | RAS-like, family 11, member B | -0.63770 | **0.64274** | 0.00005 | 0.00621 |
| Ckb | creatine kinase, brain | -0.63947 | **0.64195** | 0.00040 | 0.03491 |
| Rnd1 | Rho family GTPase 1 | -0.64186 | **0.64089** | 0.00050 | 0.04149 |
| Nat14 | N-acetyltransferase 14 | -0.64397 | **0.63995** | 0.00050 | 0.04149 |
| 2810428I15Rik | required for excision 1-B domain containing | -0.64937 | **0.63756** | 0.00010 | 0.01142 |
| Gm12892 | predicted gene 12892 | -0.65636 | **0.63448** | 0.00005 | 0.00621 |
| Ifit3 | interferon-induced protein with tetratricopeptide repeats 3 | -0.66241 | **0.63182** | 0.00045 | 0.03819 |
| Cx3cl1 | chemokine (C-X3-C motif) ligand 1 | -0.67915 | **0.62453** | 0.00015 | 0.01595 |
| Nrep | neuronal regeneration related protein | -0.70020 | **0.61549** | 0.00010 | 0.01142 |
| Wscd2 | WSC domain containing 2 | -0.70128 | **0.61503** | 0.00010 | 0.01142 |
| Nr4a1 | nuclear receptor subfamily 4, group A, member 1 | -0.70395 | **0.61389** | 0.00005 | 0.00621 |
| Glipr2 | GLI pathogenesis-related 2 | -0.70857 | **0.61193** | 0.00005 | 0.00621 |
| Relb | avian reticuloendotheliosis viral (v-rel) oncogene related B | -0.71545 | **0.60902** | 0.00005 | 0.00621 |
| Mylk2 | myosin, light polypeptide kinase 2, skeletal muscle | -0.71928 | **0.60740** | 0.00015 | 0.01595 |
| Dact3 | dishevelled-binding antagonist of beta-catenin 3 | -0.73605 | **0.60038** | 0.00005 | 0.00621 |
| Trp53i11 | transformation related protein 53 inducible protein 11 | -0.74756 | **0.59561** | 0.00005 | 0.00621 |
| Hist1h1c | histone cluster 1, H1c | -0.75072 | **0.59431** | 0.00005 | 0.00621 |
| AW112010 | expressed sequence AW112010 | -0.75645 | **0.59195** | 0.00010 | 0.01142 |
| Tcf15 | transcription factor 15 | -0.76441 | **0.58870** | 0.00015 | 0.01595 |
| Gm13889 | predicted gene 13889 | -0.77788 | **0.58322** | 0.00005 | 0.00621 |
| Ada | adenosine deaminase | -0.78105 | **0.58194** | 0.00030 | 0.02818 |
| Rn7sk | RNA, 7SK, nuclear | -0.79642 | **0.57577** | 0.00005 | 0.00621 |
| Nr4a3 | nuclear receptor subfamily 4, group A, member 3 | -0.80379 | **0.57284** | 0.00015 | 0.01595 |
| Unc119 | unc-119 lipid binding chaperone | -0.81547 | **0.56822** | 0.00005 | 0.00621 |
| Inhbb | inhibin beta-B | -0.86685 | **0.54834** | 0.00005 | 0.00621 |
| Hspa1b | heat shock protein 1B | -0.87061 | **0.54692** | 0.00030 | 0.02818 |
| Npas2 | neuronal PAS domain protein 2 | -0.87338 | **0.54587** | 0.00010 | 0.01142 |
| Slc6a2 | solute carrier family 6 (neurotransmitter transporter, noradrenalin), member 2 | -0.95912 | **0.51437** | 0.00045 | 0.03819 |
| Spn | sialophorin | -0.96711 | **0.51153** | 0.00010 | 0.01142 |
| Cyr61 | cysteine rich protein 61 | -0.97432 | **0.50898** | 0.00005 | 0.00621 |
| Fgf9 | fibroblast growth factor 9 | -1.01614 | **0.49444** | 0.00055 | 0.04523 |
| Slfn9 | schlafen 9 | -1.02316 | **0.49204** | 0.00010 | 0.01142 |
| Tnfsf10 | tumor necrosis factor (ligand) superfamily, member 10 | -1.02518 | **0.49135** | 0.00005 | 0.00621 |
| Arntl | aryl hydrocarbon receptor nuclear translocator-like | -1.03280 | **0.48876** | 0.00005 | 0.00621 |
| Irgm1 | immunity-related GTPase family M member 1 | -1.03831 | **0.48690** | 0.00010 | 0.01142 |
| Ldlr | low density lipoprotein receptor | -1.03992 | **0.48636** | 0.00005 | 0.00621 |
| Foxs1 | forkhead box S1 | -1.04255 | **0.48547** | 0.00005 | 0.00621 |
| Hspa1a | heat shock protein 1A | -1.06006 | **0.47961** | 0.00005 | 0.00621 |
| Igfbp2 | insulin-like growth factor binding protein 2 | -1.06472 | **0.47807** | 0.00020 | 0.02011 |
| Gm26917 | predicted gene, 26917 | -1.06783 | **0.47704** | 0.00005 | 0.00621 |
| Gbp5 | guanylate binding protein 5 | -1.07095 | **0.47601** | 0.00045 | 0.03819 |
| Socs1 | suppressor of cytokine signaling 1 | -1.15438 | **0.44926** | 0.00015 | 0.01595 |
| Cldn3 | claudin 3 | -1.17873 | **0.44174** | 0.00005 | 0.00621 |
| Gm12250 | predicted gene 12250 | -1.20357 | **0.43420** | 0.00005 | 0.00621 |
| Spon2 | spondin 2, extracellular matrix protein | -1.20549 | **0.43362** | 0.00005 | 0.00621 |
| Rpph1 | ribonuclease P RNA component H1 | -1.30684 | **0.40421** | 0.00005 | 0.00621 |
| Neurl3 | neuralized E3 ubiquitin protein ligase 3 | -1.35878 | **0.38991** | 0.00005 | 0.00621 |
| Atf3 | activating transcription factor 3 | -1.39387 | **0.38054** | 0.00005 | 0.00621 |
| Gzma | granzyme A | -1.40290 | **0.37817** | 0.00005 | 0.00621 |
| Apol9a | apolipoprotein L 9a | -1.46762 | **0.36158** | 0.00005 | 0.00621 |
| Nkg7 | natural killer cell group 7 sequence | -1.50521 | **0.35228** | 0.00045 | 0.03819 |
| Sele | selectin, endothelial cell | -1.56657 | **0.33761** | 0.00005 | 0.00621 |
| Ccl5 | chemokine (C-C motif) ligand 5 | -1.59601 | **0.33079** | 0.00005 | 0.00621 |
| Ido1 | indoleamine 2,3-dioxygenase 1 | -1.61053 | **0.32748** | 0.00005 | 0.00621 |
| Gzmb | granzyme B | -1.61158 | **0.32724** | 0.00005 | 0.00621 |
| Cd274 | CD274 antigen | -1.61748 | **0.32591** | 0.00005 | 0.00621 |
| Apol9b | apolipoprotein L 9b | -1.65973 | **0.31650** | 0.00005 | 0.00621 |
| Zbp1 | Z-DNA binding protein 1 | -1.68718 | **0.31053** | 0.00005 | 0.00621 |
| Apol7c | apolipoprotein L 7c | -1.75159 | **0.29698** | 0.00020 | 0.02011 |
| Tbx21 | T-box 21 | -1.88205 | **0.27130** | 0.00020 | 0.02011 |
| Il18rap | interleukin 18 receptor accessory protein | -1.96874 | **0.25548** | 0.00005 | 0.00621 |
| Cxcl9 | chemokine (C-X-C motif) ligand 9 | -2.24791 | **0.21053** | 0.00005 | 0.00621 |
| Ccdc184 | coiled-coil domain containing 184 | -2.25470 | **0.20954** | 0.00005 | 0.00621 |
| Cdh16 | cadherin 16 | -2.25993 | **0.20878** | 0.00005 | 0.00621 |
| Serpina3g | serine (or cysteine) peptidase inhibitor, clade A, member 3G | -2.37078 | **0.19334** | 0.00005 | 0.00621 |
| 2010109I03Rik | RIKEN cDNA 2010109I03 gene | -2.66587 | **0.15758** | 0.00035 | 0.03159 |
| Apoa1 | apolipoprotein A-I | -3.06859 | **0.11920** | 0.00005 | 0.00621 |
| Serpina3f | serine (or cysteine) peptidase inhibitor, clade A, member 3F | -4.08432 | **0.05895** | 0.00005 | 0.00621 |
| Ubd | ubiquitin D | -4.29165 | **0.05106** | 0.00005 | 0.00621 |
| 6430584L05Rik | RIKEN cDNA 6430584L05 gene | -4.31371 | **0.05029** | 0.00005 | 0.00621 |
| Gm6637 | predicted gene 6637 | -4.92523 | **0.03291** | 0.00005 | 0.00621 |
